# Integrated Single-Cell Genotyping and Chromatin Accessibility Charts *JAK2^V617F^* Human Hematopoietic Differentiation

**DOI:** 10.1101/2022.05.11.491515

**Authors:** Robert M. Myers, Franco Izzo, Sanjay Kottapalli, Tamara Prieto, Andrew Dunbar, Robert L. Bowman, Eleni P. Mimitou, Maximilian Stahl, Sebastian El Ghaity-Beckley, JoAnn Arandela, Ramya Raviram, Saravanan Ganesan, Levan Mekerishvili, Ronald Hoffman, Ronan Chaligné, Omar Abdel-Wahab, Peter Smibert, Bridget Marcellino, Ross L. Levine, Dan A. Landau

## Abstract

In normal somatic tissue differentiation, changes in chromatin accessibility govern priming and commitment of precursors towards cellular fates. In turn, somatic mutations can disrupt differentiation topologies leading to abnormal clonal outgrowth. However, defining the impact of somatic mutations on the epigenome in human samples is challenging due to admixed mutated and wildtype cells. To chart how somatic mutations disrupt epigenetic landscapes in human clonal outgrowths, we developed Genotyping of Targeted loci with single-cell Chromatin Accessibility (GoT-ChA). This high-throughput, broadly accessible platform links genotypes to chromatin accessibility at single-cell resolution, across thousands of cells within a single assay. We applied GoT-ChA to CD34^+^ cells from myeloproliferative neoplasm (MPN) patients with *JAK2^V617F^*-mutated hematopoiesis, where the *JAK2* mutation is known to perturb hematopoietic differentiation. Differential accessibility analysis between wildtype and *JAK2^V617F^* mutant progenitors revealed both cell-intrinsic and cell state-specific shifts within mutant hematopoietic precursors. An early subset of mutant hematopoietic stem and progenitor cells (HSPCs) exhibited a cell-intrinsic pro-inflammatory signature characterized by increased NF-κB and JUN/FOS transcription factor motif accessibility. In addition, mutant HSPCs showed increased myeloid/erythroid epigenetic priming, preceding increased erythroid and megakaryocytic cellular output. Erythroid progenitors displayed aberrant regulation of the γ-globin locus, providing an intrinsic epigenetic basis for the dysregulated fetal hemoglobin expression observed in MPNs. In contrast, megakaryocytic progenitors exhibited a more specialized inflammatory chromatin landscape relative to early HSPCs, with increased accessibility of pro-fibrotic JUN/FOS transcription factors. Notably, analysis of myelofibrosis patients treated with JAK inhibitors revealed an overall loss of mutant-specific phenotypes without modifying clonal burden, consistent with clinical responses. Finally, expansion of the multi-modality capability of GoT-ChA to integrate mitochondrial genome profiling and cell surface protein expression measurement enabled genotyping imputation and discovery of aberrant cellular phenotypes. Collectively, we show that the *JAK2^V617F^* mutation leads to epigenetic rewiring in a cell-intrinsic and cell type-specific manner. We envision that GoT-ChA will thus serve as a foundation for broad future explorations to uncover the critical link between mutated somatic genotypes and epigenetic alterations across clonal populations in malignant and non-malignant contexts.

## INTRODUCTION

Differentiation topologies are critical for homeostasis across human tissues and are maintained through tightly coordinated epigenetic regulation. For example, in the hematopoietic system, coordinated epigenetic changes are reflected in the chromatin accessibility landscape and determine hematopoietic and stem progenitor cells (HSPCs) commitment towards specific downstream cellular fates^1^. Recent advances in single cell mapping of chromatin accessibility revealed that key transcription factors promote variability in the accessibility of regulatory elements, underlying the epigenetic heterogeneity across progenitor cells^2^. Importantly, during hematopoiesis, shifts in chromatin accessibility at regulatory regions precede transcriptional changes during differentiation^3^, a process defined as epigenetic priming^4^.

Somatic driver mutations within HSPCs result in clonal expansions and the reshaping of differentiation landscapes. Altered HSPC differentiation landscapes range from subtle differentiation biases in clonal hematopoiesis^5,6^, through skewed accumulation of mature blood cells in myeloproliferative neoplasia (MPN)^7,8^, to impaired differentiation in leukemogenic transformation^9,10^. The recurrent V617F hotspot mutation in the Janus Kinase 2 (*JAK2*) gene serves as a prototypical example, associated both with clonal hematopoiesis^11–14^ as well as overt differentiation skews in polycythemia vera (PV) and essential thrombocythemia (ET), and ultimately leading to bone marrow failure in myelofibrosis (MF)^15–18^. The JAK2^V617F^ protein is constitutively active and phosphorylates downstream signal transducers and activators of transcription (STATs)^16,17^, which promote an inflammatory phenotype^19–21^ and result in cytokine-independent expansion of the erythroid-megakaryocytic lineage^7,17,22,23^. However, in humans, mutated and wildtype cells are often admixed without clear distinguishing cell surface markers amenable to fluorescence-activated cell sorting (FACS) enrichment, limiting bulk or single-cell chromatin profiling. Moreover, inter-individual comparisons are limited by clinical confounders, as well as potential cell-extrinsic effects due to variation of microenvironmental cues across patients^24–26^. Thus, the epigenetic underpinning of *JAK2^V617F^* differentiation topographies in human disease remains largely unknown.

To overcome the challenge of resolving admixed wildtype and mutated cells in primary human samples, single-cell multi-omics methods have been developed to link genotypes with transcriptional^24,25,27–33^ or cell surface protein^34,35^ profiles. TARGET-seq, for example, is a plate-based technique that performs simultaneous single-cell RNA-seq (scRNA-seq) with genotyping via targeted amplification of both complementary DNA (cDNA) and genomic DNA (gDNA), achieving up to 98% detection rate of multiple mutations in hundreds of single cells at a time^25,27^. Alternative droplet- or nanowell-based techniques allow for higher throughput with simultaneous scRNA-seq and genotyping of the actively transcribed genes containing the loci of interest^28,29,31,32^. However, these droplet-based genotyping methods rely solely on captured mRNA transcripts, which results in a limiting dependency on both the expression level of the targeted gene and the distance of the mutated locus from transcript end. Indeed, these features have been shown to severely impact the ability of transcription-based genotyping for lowly expressed genes, such as *JAK2*^29,31^. Finally, existing methods are unable to jointly capture genotyping and chromatin accessibility, a requirement for the study of the impact of somatic mutations on the epigenetic landscape of clonal HSPCs.

Here, we developed <u>G</u>enotyping <u>o</u>f <u>T</u>argeted loci with single-cell <u>C</u>hromatin <u>A</u>ccessibility (GoT-ChA), for droplet-based high-throughput simultaneous capture of genotyping and chromatin accessibility from the same single cell. This approach allows for high resolution intra-sample comparisons of mutated versus wildtype chromatin accessibility profiles. As wildtype cells within the same sample provide the ideal comparator to mutant cells, GoT-ChA bypasses clinical confounders in inter-individual comparisons in human studies, as well as allowing to decouple cell-intrinsic from microenvironment effects. Unlike RNA-based genotyping methods, single cell genotyping with GoT-ChA is based on targeted amplification of gDNA. Thus, GoT-ChA obviates both the dependency on expression level of the gene containing the target locus, as well as the dependency on the mutation position relative to the expressed transcript end, radically expanding the scope of genetic alterations that can be assayed with high-throughput single cell profiling. Furthermore, we integrated GoT-ChA with copy number variation (CNV) inference^36^ as well as mitochondrial genome and cell surface protein capture in single cells^37,38^, delivering a highly multi-modal single-cell platform.

When applied to human PV and MF samples targeting the *JAK2^V617^* locus, GoT-ChA drastically increased the rate of genotyped cells relative to RNA-based methods^29,31,32^. While untreated patients showed erythroid and megakaryocytic differentiation biases consistent with the disease phenotype^39^, patients treated with JAK2 inhibitors show a more homogeneous distribution of mutant cells across the differentiation topology, suggesting that JAK inhibition abrogates differential hematopoietic contributions of *JAK2^V617F^* without eliminating disease-initiating HSPCs, in line with clinical observations^40–42^. Intra-individual and intra-cluster comparisons show that cell-intrinsic pro-inflammatory epigenetic profiles are already present in a subset of the earliest *JAK2^V617F^* mutant HSPCs, while increased accessibility of transcription factor motifs involved in erythroid differentiation consistent with epigenetic priming is observed across HSPCs. In *JAK2^V617F^* committed erythroid progenitors, we observed increased accessibility of the *HBG1* gene, a component of fetal hemoglobin (HbF), while megakaryocyte progenitors showed increased motif accessibility of pro-fibrotic JUN/FOS^43–45^ transcription factors. These observations underscore intrinsic and cell type-specific effects of the *JAK2^V617F^* mutation in human hematopoiesis and demonstrate the ability of GoT-ChA to resolve clonal admixtures and to provide genotype to epigenome mapping of clonal outgrowths in primary human samples.

## RESULTS

### Droplet-based high-throughput simultaneous capture of genotypes and chromatin accessibility in single cells

To integrate genotyping into scATAC-seq, we modified the broadly utilized 10x Genomics platform by adding two custom primers (GoT-ChA primers) to the cell barcoding PCR reaction mixture prior to loading the microfluidics chip for droplet generation (**Fig. 1a**). These primers are designed to flank a specific region of interest, with one primer containing a Read 1 Nextera (R1N) handle (**Extended Fig. 1a-b**). During in-droplet PCR, the GoT-ChA primers bind to and amplify the genomic region of interest, generating an amplicon that is complementary to the 10x scATAC-seq Gel Bead oligonucleotides, allowing for cell barcoding and further amplification. In this manner, both tagmented gDNA fragments and GoT-ChA amplicons simultaneously obtain the unique cell barcode sequence and the P5 Illumina sequencing handle prior to breakage of the single-cell emulsion (**Fig. 1a**). Of note, tagmentation of the targeted site is not required for genotype capture, as target amplification with GoT-ChA primers results in amplicons already containing the R1N capture sequence. Moreover, while in-droplet amplification of tagmented genomic fragments is linear, GoT-ChA amplicons are exponentially amplified by design, resulting in an increased relative abundance of genotyping amplicons. Then, 10% of the total product is used for genotyping library construction, while the remainder is utilized for the scATAC-seq library. During genotyping library construction, GoT-ChA fragments are specifically amplified using a hemi-nested PCR strategy and a biotin-streptavidin pulldown prior to sample indexing (**Extended Data Fig. 1c**). Both the scATAC-seq and GoT-ChA libraries can then be sequenced together, with chromatin accessibility and genotype information linked via shared cell barcodes. To encourage the broad implementation of GoT-ChA, we developed an R package (Gotcha R package, **see code availability section**) that provides start-to-end processing of GoT-ChA data and its integration with the corresponding scATAC-seq profiles.

**Fig. 1.**
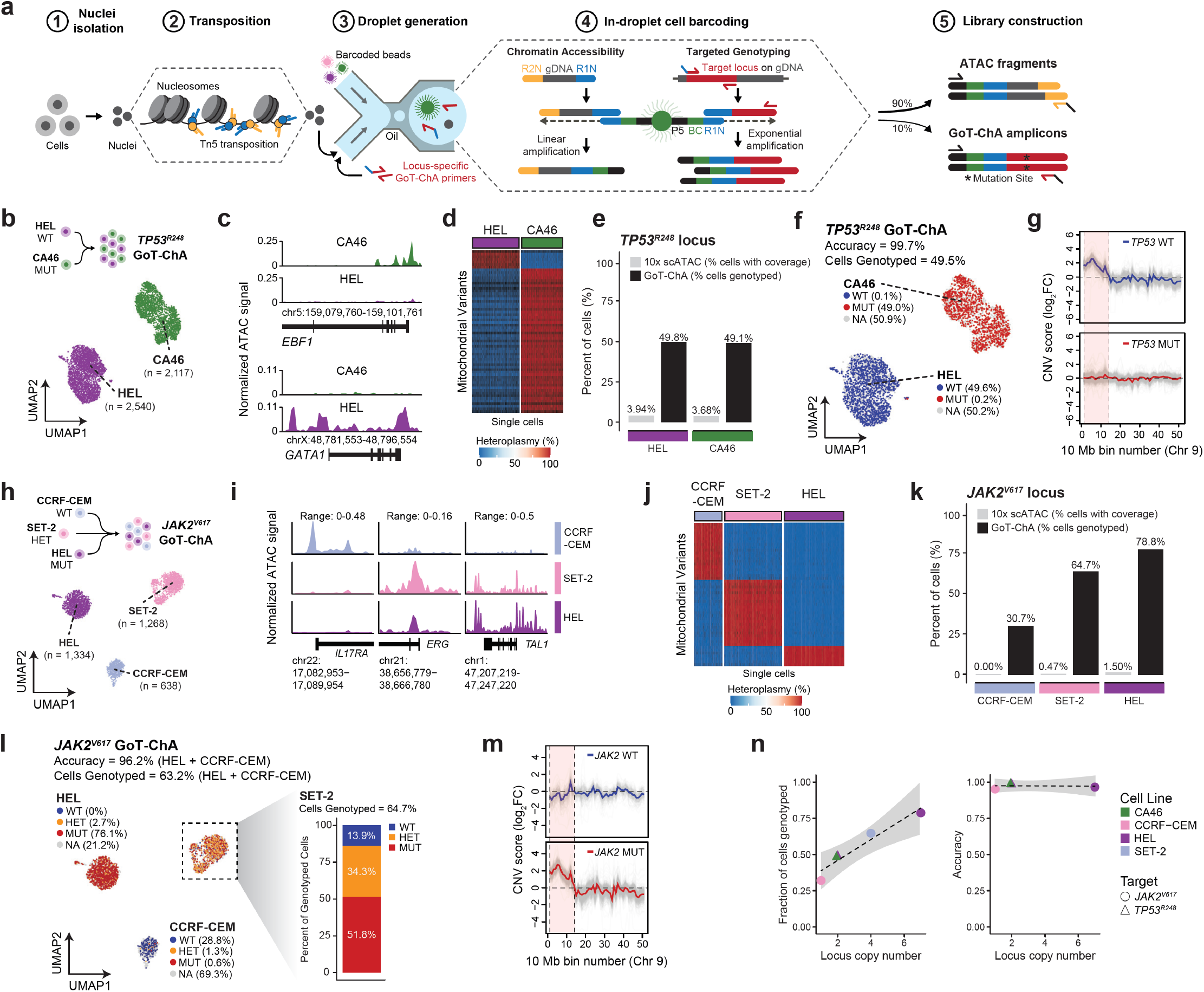
GoT-ChA allows for accurate single-cell genotyping of somatic mutations together with chromatin accessibility information. **a**, Schematic representation of GoT-ChA workflow. P5, Illumina sequencing handle; BC, unique cell barcode; R1N, Read 1 Nextera adapter; gDNA, genomic DNA; R2N, Read 2 Nextera adapter. **b**, Schematic representation of the *TP53^R248^* mixing study (upper panel) and accessibility-based uniform manifold approximation and projection (UMAP) for HEL (n = 2,540 cells; *TP53^WT/WT^*; purple) and CA46 (n = 2,117 cells; *TP53^MUT/MUT^*; green; bottom panel). **c**, Chromatin accessibility coverage of marker genes (FDR < 0.05 and log_2_FC > 1.25; Wilcoxon rank sum test followed by Benjamini-Hochberg correction), agnostic to genotyping information, used for cell line identity assignments. ATAC signal was normalized by the number of reads covering transcriptional start sites (TSS). **d**, Heatmap showing heteroplasmy for mutually exclusive mitochondrial variants detected in the scATAC-seq data for either HEL or CA46 cell lines. Minimum coverage of 10 reads mapping to the variant site was required. **e**, Detected coverage for the *TP53^R248^* locus of interest for either HEL or CA46 cell lines illustrating improved genotyping capture with GoT-ChA versus standard 10x scATAC. Percent of cells reported for the 10x scATAC libraries was defined as the percent of cells with at least one read mapping to the assayed locus. Percent of cells for the GoT-ChA libraries was defined as successful genotyping post processing (see materials and methods). **f**, UMAP colored by GoT-ChA genotype classifications of CA46 (n = 2,117 cells) and HEL (n = 2,540 cells) assigned as wildtype (WT, blue), mutant (MUT, red), or not assignable (NA, grey) cells. The percentage of cells assigned to each genotype per cell line, as well as accuracy and overall percent of cells genotyped are shown. **g**, Inferred copy number variation (CNV) scores from the scATAC-seq data (see materials and methods) in either *TP53^R248^* wildtype or mutant cells as defined by GoT-ChA mixing study relative to a healthy control (see materials and methods), verifying the concordance between known amplification of the region including the *JAK2* gene on chromosome 9 in HEL cells, which are wildtype for *TP53^R248^*. **h**, Schematic representation of the *JAK2^V617^* mixing study (top panel) and chromatin accessibility-based UMAP for HEL (n = 1,334 cells; *JAK2^MUT/MUT^*; purple), CCRF-CEM (n = 638 cells; *JAK2^WT/WT^*; light blue) and SET-2 (n = 1,268 cells; *JAK2^WT/MUT^*; pink) cell lines (bottom panel). **i**, Chromatin accessibility coverage of marker genes (FDR < 0.05, log_2_FC > 1.25), agnostic to genotyping information used for cell line identity assignments. Wilcoxon rank sum test followed by Benjamini-Hochberg correction. **j**, Heatmap showing heteroplasmy of mutually exclusive mitochondrial variants detected in the scATAC-seq data for HEL, CCRF-CEM and SET-2 cell lines. Minimum coverage of 10 reads mapping to the variant site was required. **k**, Detected coverage for the locus of interest per cell line illustrating improved genotyping capture with GoT-ChA versus standard 10x scATAC. Percent of cells reported for 10x scATAC-seq and GoT-ChA are as defined in panel **e**. **l**, Chromatin accessibility-based UMAP for HEL (n = 1,334 cells), SET-2 (n = 1,268 cells) and CCRF-CEM (n = 638 cells) colored by GoT-ChA genotype classifications of homozygous wildtype (WT, blue), homozygous mutant (MUT, red), heterozygous (HET, yellow), and not assignable (NA, grey) cells (left panel). Percentage of cells assigned to each genotype and overall accuracy and percent of genotyped cells is reported for the HEL and CCRF-CEM homozygous cell lines. Genotyping results for the SET-2 cell line indicating multi-allelic capture by GoT-ChA genotyping in a subset of cells, allowing for accurate heterozygous classification in 34.3% of genotyped cells (right panel). We note that mutated and wildtype annotation likely represents incomplete capture of all alleles, and the observed percentages correspond to the 3:1 allelic ratio of mutated:wildtype alleles present in SET-2 cells, as verified by Sanger sequencing (see **Extended Data 2k**). **m**, CNV scores inferred from the scATAC-seq data (see materials and methods) comparing homozygous wildtype and mutant cells in the *JAK2^V617^* GoT-ChA mixing study relative to a healthy control (see materials and methods), illustrating concordance between known amplification of the region including the *JAK2* gene on chromosome 9 in HEL cells, which are mutant for *JAK2^V617^*. **n**, GoT-ChA quality metrics in relation to target locus copy number: fraction of cells genotyped (left panel) positively correlates with locus copy number, while genotyping accuracy (right panel) remains above 95% irrespective of locus copy number.

To test the ability of GoT-ChA to accurately capture genotype along with chromatin accessibility information, we performed a cell line mixing study. Two human cell lines of discrete cell types (CA46, a B lymphocyte cell line; and HEL, an erythroblast cell line) and differing genotypes for the *TP53^R248^* locus (CA46, *TP53^R248Q^* homozygous mutant; HEL, *TP53^R248^* homozygous wildtype; **Extended Data Fig. 2a**) were mixed at a 1:1 ratio and profiled with GoT-ChA (**Fig. 1b, upper panel**). Using chromatin accessibility information alone, cells clustered into two distinct populations (**Fig. 1b, bottom panel**). The two clusters were readily annotated as the expected mixed cell lines via chromatin accessibility signals at key marker genes (**Fig. 1c**, **Extended Data Fig. 2b**), as well as mutually exclusive mitochondrial variants (**Fig. 1d**). Fragment size distribution, total number of fragments per cell, and enrichment of fragments mapping to transcriptional start sites (TSS) all reflect high quality scATAC-seq data, unaffected by the inclusion of GoT-ChA primers during droplet generation (**Extended Data Fig. 2c-e**).

Analysis of the matching unprocessed genotyping data showed the presence of two distinct modes in the distribution of genotyping reads per cell. We reasoned that these distributions reflect cells for which genotyping was successfully captured versus cells displaying background noise from the genotyping library. To address this aspect, we developed a computational framework (**Extended Data Fig. 2f**) using kernel density estimation (KDE) to define the boundaries between background noise and genotyping signal (**Extended Data Fig. 2g**), followed by clustering and genotype assignment based on the z-scores of genotyping read counts (**Extended Data Fig. 2h**). This approach was orthogonally validated using an alternative framework leveraging recent work^46^ in single-cell genomics that proposed estimating the background noise with empty droplets (**Extended Data Fig. 2i-j**; **see materials and methods**; both approaches for noise correction and genotype calling are included in the Gotcha R package). While less than 4% of cells had at least one scATAC-seq read covering the *TP53^R248^* locus of interest in either cell line, GoT-ChA resulted in high confidence genotyping for 49.8% and 49.1% of HEL and CA46 cells, respectively (**Fig. 1e**). By correcting for background noise, we achieved genotyping of the *TP53^R248^* locus in 49.5% of all cells with an accuracy of 99.7% (**Fig. 1f**). As an additional orthogonal validation of our genotyping, we compared wildtype and mutant cell CNV scores^36^ inferred from the chromatin accessibility profiles (**see materials and methods**) across chromosome 9, for which the *TP53^R248^* wildtype HEL cell line carries an amplification^47–49^. Consistent with our observed genotyping, wildtype cells as defined by GoT-ChA genotyping exhibited an increased CNV score relative to mutant cells, further confirming successful single-cell genotyping (**Fig. 1g**).

We further tested GoT-ChA targeting a different genomic locus, the *JAK2^V617^* hotspot, with a separate mixing study to directly address heterozygous genotyping. Three human cell lines (HEL, an erythroblast cell line; SET-2, a megakaryoblast cell line; CCRF-CEM, a T lymphoblast cell line) with discrete *JAK2^V617^* genotypes (HEL, *JAK2^V617F^* homozygous mutant; SET-2, *JAK2^V617F^* heterozygous; CCRF-CEM, *JAK2^V617^* homozygous wildtype; **Extended Data Fig. 2k**) were mixed at a 1:1:1 ratio and profiled with GoT-ChA. As with the homozygous mixing study targeting *TP53^R248^* shown above, clustering agnostic to genotyping utilizing only the scATAC-seq data resulted in discrete populations of cells (**Fig. 1h**) that were annotated via differential chromatin accessibility signals at key marker genes (**Fig. 1i**; **Extended Data Fig. 2l**) and unique mitochondrial variants (**Fig. 1j**, while again confirming unaffected quality of the scATAC-seq data (**Extended Data Fig. 2m, n**). The genotyping data was processed as described above, using KDE applied to log-read distributions to define the boundaries between background noise and genotyping signal for wildtype and mutant reads (**Extended Data Fig. 2o**), followed by clustering on z-score read counts (**Extended Data Fig. 2p**). While less than 2% of cells had at least one scATAC-seq read covering the *JAK2^V617^* locus, genotyping by GoT-ChA captured 30.7% of CCRF-CEM, 64.7% of SET-2 and 78.8% of HEL cells (**Fig. 1k**). In the two homozygous cell lines (wildtype CCRF-CEM and mutant HEL), genotyping of the *JAK2^V617^* locus was achieved in 63.2% of cells with an accuracy of 96.2% (**Fig. 1l, left panel**). In the SET-2 heterozygous cell line, 64.7% of cells were successfully genotyped; 34.3% of genotyped cells were correctly identified as heterozygous, confirming bi-allelic capture, with the remaining genotyped cells split between homozygous wildtype and mutant assignments, suggesting incomplete capture of the heterozygous genotype (**Fig. 1l, right panel,** note that SET-2 cells have a 3:1 ratio of mutated:wildtype alleles (**Extended Data Fig. 2k**), likely underlying the higher fraction of mutated versus wildtype calls in non-heterozygous genotype classification). CNV scores for homozygous wildtype and mutant cells along the chromosome 9 amplification present in the *JAK2^V617F^* homozygous mutant HEL cell line^47–49^ orthogonally validated successful genotyping of the *JAK2^V617^* locus (**Fig. 1m**). We note that variation in genotyping efficiency across cell lines is likely related to the copy number variation for the *JAK2* locus^47,50,51^. Indeed, the proportion of genotyped cells positively correlated with the copy number of the *JAK2* locus (**Fig. 1n, left panel**, Pearson correlation; *P* = 0.011; R^2^ = 0.91; F-test), suggesting that additional copies of the target locus improve the genotype capture rate. Importantly, genotyping accuracy remained consistent regardless of target copy number differences (**Fig. 1n, right panel**).

Altogether, these data demonstrate that GoT-ChA allows for high-throughput simultaneous capture of genotypes and chromatin accessibility profiles in single cells, with high accuracy and cell recovery independent of expression level and genomic localization of the targeted region.

### GoT-ChA of primary human *JAK2^V617F^* myelofibrosis reveals cell type-specific mutant cell predominance in erythroid and megakaryocytic progenitors

The *JAK2^V617F^* mutation has a central role in the pathogenesis of myeloproliferative neoplasms^7,15,18,22^. We sought to explore how this mutation disrupts the regulatory chromatin landscape that determines cell-fate decisions of HSPCs. To address this question, we applied GoT-ChA to CD34^+^ sorted progenitor cells from seven patients with *JAK2^V617F^-mutated* MF with no additional mutations (**Extended Data Table 1**), who had either not been treated with JAK inhibition or were being treated with ruxolitinib, a JAK1/2 inhibitor, or fedratinib, a JAK2-specific inhibitor at the time of sample collection (**Extended Data Table 1**). The quality of the scATAC-seq data was not affected by removal of a small portion of the sample for GoT-ChA genotyping library construction (**Extended Data Fig. 3a-c**). We then performed cell clustering in a manner agnostic to genotyping information, based solely on chromatin accessibility (**Fig. 2a**). Reciprocal latent semantic indexing^52^ applied to the binarized genomic bin by cell matrix was used for sample integration to correct for patient-specific batch effects (**Fig. 2b; Extended Data Fig. 3d; see materials and methods**). Individual cluster identities were assigned using gene accessibility scores of key marker genes for hematopoietic lineages (**Fig. 2c; Extended Data Fig. 3e**) and confirmed via transcription factor footprinting (**Fig. 2d**) and peak calling (**Extended Data Fig. 3f**), followed by differential enrichment of marker peaks (**Extended Data Fig. 3g**). We identified the expected progenitor subtypes, along with a population of mature monocytic cells characterized by CD14 expression and lack of CD34 expression, often observed in CD34^+^ sorting of human bone marrow^53^. The presence of two distinct HSPC subclusters, early (HSPC1) and late (HSPC2), was validated via gene accessibility scores of multiple markers of hematopoietic stem cells (**Extended Data Fig. 3h**). Cluster assignments were further supported via orthogonal annotation utilizing cell mapping through a novel multiomic bridge integration approach^54^ (**Extended Data Fig. 3i; see materials and methods**).

**Fig. 2.**
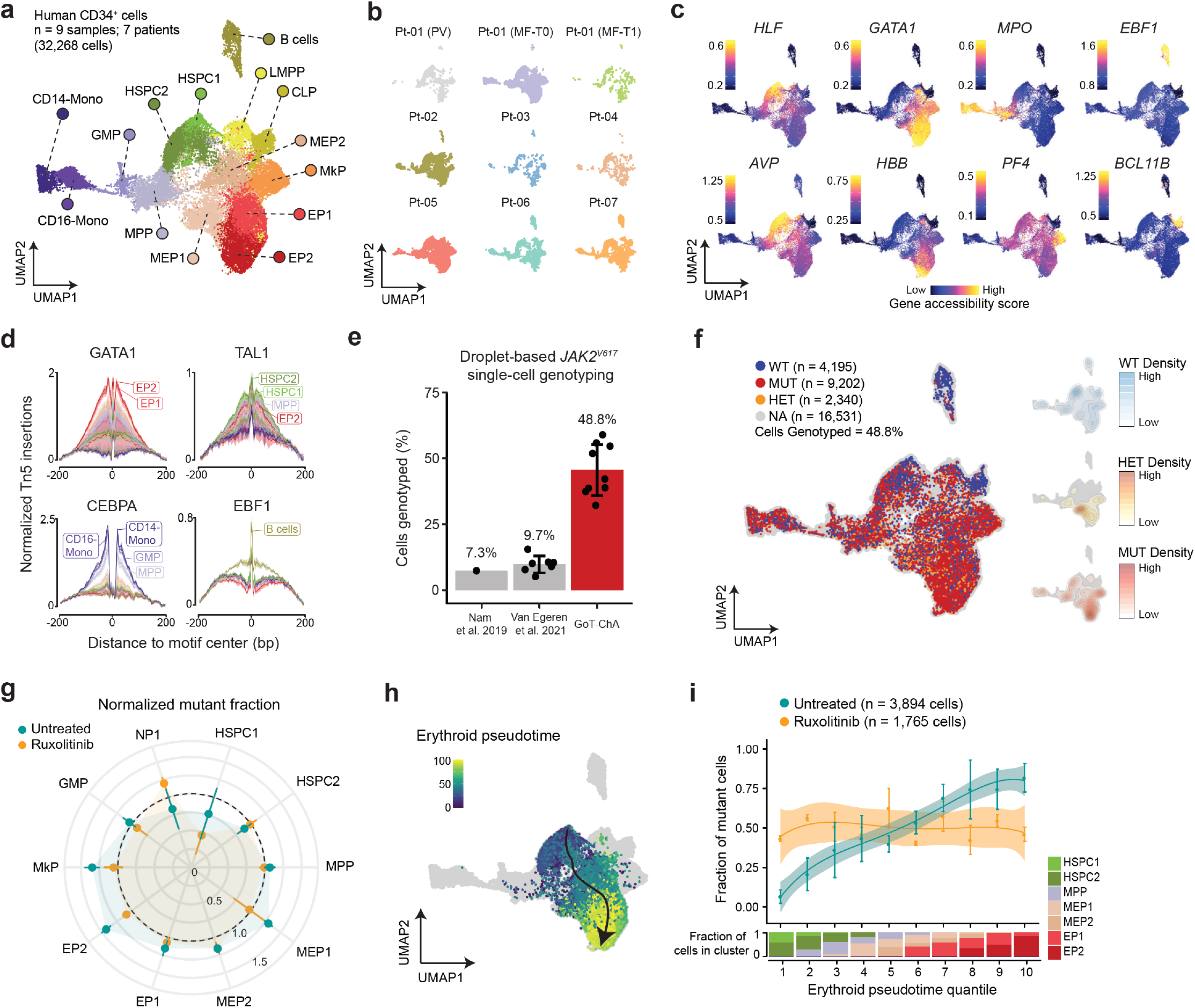
GoT-ChA on human *JAK2^V617F^*-mutated myelofibrosis samples reveals myeloid differentiation bias that is abrogated with ruxolitinib treatment. **a**, Integrated chromatin accessibility uniform manifold approximation and projection (UMAP) after reciprocal latent semantic indexing (LSI) integration (see materials and methods) from CD34^+^ sorted patient samples (n = 9 samples, 7 patients, 32,268 cells) illustrating the expected progenitor populations in early hematopoiesis. HSPC = hematopoietic stem and progenitor cells; MPP = multi-potent progenitors; GMP = granulocyte monocyte progenitors; MEP = megakaryocyte-erythrocyte progenitors; EP = erythroid progenitors; MkP = megakaryocytic progenitors; LMPP = lymphoid-myeloid pluripotent progenitors; CLP = common lymphoid progenitors; CD16-Mono = CD16^+^ monocyte progenitors; CD14-Mono = CD14^+^ monocytes. **b**, Integrated UMAP plotted for each patient sample, illustrating consistent distribution of cells after reciprocal LSI integration (see materials and methods). **c**, UMAPs showing the gene accessibility scores of canonical marker genes (FDR < 0.05 and log_2_FC > 1; Wilcoxon rank sum test followed by Benjamini-Hochberg correction) used to identify the individual progenitor clusters within the integrated UMAP. **d**, Transcription factor footprinting of canonical transcription factors for progenitor clusters, further confirming cluster identity assignments. **e**, Comparison of *JAK2^V617^* genotyping efficiency across recent publications applying single-cell droplet-based technologies for genotyping. Bars represent mean values, each point represents an individual patient sample, and error bars indicate standard deviation. **f**, Integrated UMAP colored by GoT-ChA-assigned *JAK2^V617^* genotypes as homozygous wildtype (WT; n = 4,195 cells; blue), homozygous mutant (MUT; n = 9,202 cells; red) or heterozygous (HET; n = 2,340 cells; yellow) and not assignable (NA; n = 16,531 cells; grey). **g**, Radar plot showing the mutant fraction for each cluster, relative to the fraction of mutant cells observed for the entire sample, of either untreated (green) or ruxolitinib-treated (yellow) patient samples across myeloid and HSPC/MPP clusters. **h**, UMAP depicting semi-supervised pseudotime estimation for the erythroid differentiation trajectory, with the HSPC1 cluster as initial state and the EP2 cluster as final state. **i**, Fraction of mutant cells along erythroid pseudotime for untreated (n = 3,894 cells) or ruxolitinib-treated (n = 1,765 cells) samples. Erythroid pseudotime was divided in 10 quantiles; each point represents the mean fraction of mutant cells, error bars indicate standard error, lines indicate the fit and shadowed areas represent the 95% confidence interval of the generalized additive model (upper panel). The fraction of cells belonging to the cluster specified by color (HSPC = hematopoietic stem and progenitor cell; MPP = multipotent progenitor; MEP = megakaryocyte erythrocyte progenitor; EP = erythroid progenitor) within the indicated pseudotime quantile is shown (bottom panel).

Analysis of the GoT-ChA genotyping data (**Extended Data Fig. 4a**) resulted in *JAK2^V617^* genotyping information for 15,737 out of 32,268 total cells (48.8% ± 9.64%; mean ± standard deviation across samples), compared to 7-10% of cells in previously reported droplet-based scRNA-seq cDNA-based genotyping^29,31^ (**Fig. 2e**), demonstrating the increased ability of GoT-ChA to genotype loci with relatively low expression in single cells. Pseudo-bulked GoT-ChA variant allele frequencies (VAFs) were positively correlated with the reported VAFs from clinical targeted panels (*P* = 0.036; Rho = 0.84; Spearman correlation; **Extended Data Fig. 4b**). We noted variability in genotyping efficiency across cell types (range: 23.7% to 66.6%), which may be related to locus accessibility (**Extended Data Fig. 4c**). Nonetheless, by design, GoT-ChA is not strictly dependent on accessibility, allowing genotyping across cell types despite minimal locus accessibility (**Extended Data Fig. 4d**).

Projection of genotypes onto the cell differentiation map demonstrated intermingling of *JAK2* wildtype and mutant cells throughout HSPCs and myeloid progenitor clusters, while the common lymphoid progenitors (CLPs) and B cell clusters were mainly comprised of wildtype cells (**Fig. 2f, left panel; Extended Data Fig. 4e**), consistent with previous studies of hematopoietic colonies/subsets from MPN patients^23,55^. The intermingling of mutated and wildtype progenitors reinforced the inability of chromatin accessibility profiles alone to distinguish wildtype from mutant cells, highlighting the need for single-cell multi-omics for genotype-epigenome inferences.

Nonetheless, projection of genotype densities onto the differentiation map suggested an uneven distribution of mutant cells across cell subtypes (**Fig. 2f, right panel**). Indeed, while HSPCs and committed myeloid progenitor clusters were composed of an admixture of wildtype and *JAK2^V617F^* mutant cells, their frequencies varied greatly. The normalized mutant fraction was increased in the late erythroid progenitor cluster (EP2), as well as in the megakaryocytic progenitor (MkP) cluster (**Fig. 2g; Extended Data Fig. 4f**). In contrast, patients treated with the JAK1/JAK2 inhibitor ruxolitinib displayed a more homogeneous distribution of mutant cells across cell types (**Fig. 2g;** see **Extended Data Fig. 4f** for per sample analysis), suggesting that JAK inhibition alters the relative contribution of *JAK2^V617F^* to these cellular lineages, but does not eliminate the mutated clone^40–42^. Treatment with fedratinib, a JAK2-specific inhibitor, showed a similar distribution to ruxolitinib across cell types (**Extended Data Fig. 4g**). Consistently, pseudotemporal ordering of cells along the erythroid differentiation trajectory (**Fig. 2h**) showed a steady increase in the fraction of mutant cells in untreated patients, while ruxolitinib treatment resulted in a more uniform distribution of mutant cells along erythroid differentiation (**Fig. 2i**). We observed a similar pattern of mutant cell fraction increase with MkP differentiation, and to a lesser extent in the monocyte differentiation trajectory (**Extended Data Fig. 4h-i**). Thus, high resolution mapping of the *JAK2^V617F^* mutation across human hematopoietic differentiation demonstrated a progenitor-specific mutant cell predominance in myelofibrosis that is eliminated by therapeutic JAK2 inhibition.

### *JAK2^V617F^* HSPCs show a cell-intrinsic pro-inflammatory phenotype and myeloid/erythroid epigenetic priming

Inflammatory disruption of the bone marrow microenvironment has been extensively documented in myelofibrosis^19–21,39,56–59^. Indeed, secretion of inflammatory cytokines is a central feature of MPN pathophysiology and has been shown to provide a supportive niche for the expansion of mutant clones in various disease states, including MPNs and clonal hematopoiesis^33,60–63^. However, defining how wildtype and *JAK2^V617F^* mutant early progenitors differ regarding cell-intrinsic epigenetic profiles in human myelofibrosis remains unknown due to the inability to directly compare mutated and wildtype cells within primary human samples.

To delineate the effects of*JAK2^V617F^* on chromatin accessibility, we first compared gene accessibility scores (as the number of cells profiled varied between samples, linear mixture model (LMM) was used with patient sample identity explicitly modeled as random effect to account for inter-patient variability, followed by likelihood ratio test; **see materials and methods**) between wildtype and *JAK2^V617F^-mutated* cells within the HSPC1 cluster (**Fig. 3a; Extended Data Table 2**) as a proxy for gene expression. These data revealed a cell-intrinsic pro-inflammatory phenotype in *JAK2^V617F^* mutant HSPCs, with increased gene accessibility of critical inflammatory genes (**Fig. 3a-b**). Pro-inflammatory genes more accessible in *JAK2^V617F^* HSPC1 cells included members of the tumor necrosis factor (TNF) family, such as the receptor *TNFRS9* (CD137) and its ligand *TNFS9*, as well as the cytokine CD70, which acts as ligand of the *TNFRSF27* (CD27) receptor. This suggests an increase in levels of both ligands and their corresponding receptors in mutant HSPCs, which could result in increased downstream activation of NF-κB signaling^64^. In addition, both colony stimulating factors 1 and 2 (*CSF1* and *CSF2*) show increased accessibility in *JAK2^V617F^* HSPC1 cells. CSF1 can promote differentiation of HSPCs towards the myeloid lineage^65^ and CSF2 has been shown to be required for differentiation of precursor cells towards the myeloid/erythroid fate^66^. We also observed increased accessibility of the *CCL2* gene, a key cytokine mediating monocyte recruitment to inflammatory sites^67^, which has been reported to be increased in MF^68^.

**Fig. 3.**
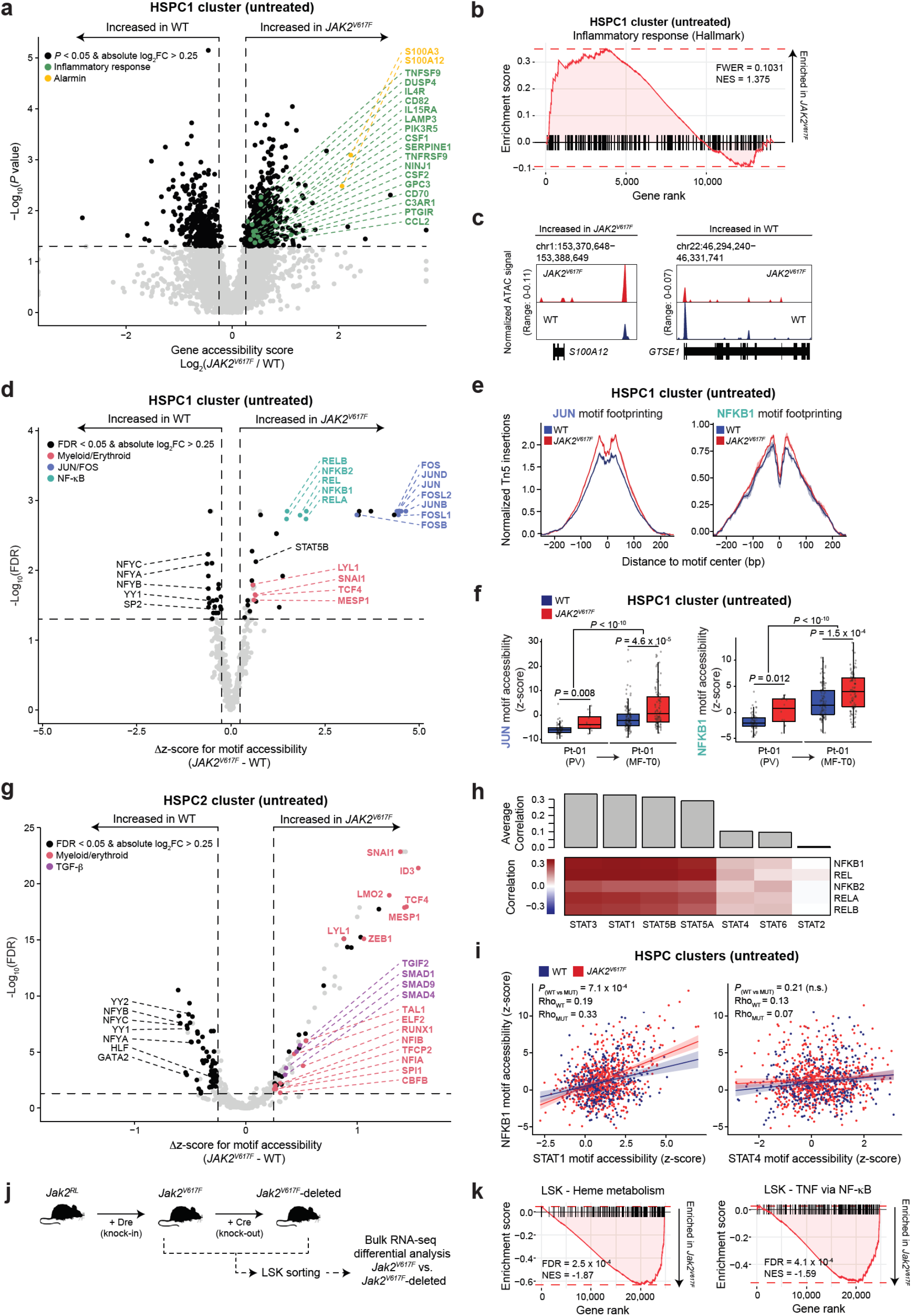
*JAK2^V617F^* mutant HSPCs exhibit intrinsic pro-inflammatory and myeloid-biased epigenetic priming that is reduced with JAK inhibition. **a**, Volcano plot illustrating differential gene accessibility scores between wildtype (n = 128 cells) and mutant (n = 93 cells) cells within the HSPC1 cluster of untreated MF patients (n = 4; excluding PV sample). Horizontal dotted line represents *P* = 0.05; vertical dotted lines represent absolute log_2_FC > 0.25. Genes involved in the inflammatory response pathway (Hallmark M5932) are highlighted in green. Alarmin genes are highlighted in yellow. Linear mixture model (LMM) modeling patient sample as a random effect for accounting for inter-patient variability followed by likelihood ratio test (see materials and methods). **b**, Pre-ranked gene set enrichment of genes within the Inflammatory response pathway for wildtype vs *JAK2^V617F^* HSPC1 gene accessibility scores (Bonferroni correction; FWER = family wise error rate; NES = normalized enrichment score; Hallmark pathway M5932). **c**, Chromatin accessibility coverage tracks of two differentially accessible (*P* < 0.05 and absolute log_2_FC > 0.25) genes between wildtype (blue) and mutant cells (red). **d**, Volcano plot of the differentially accessible transcription factor motifs (FDR < 0.05 and absolute Δz-score > 0.25; LMM followed by likelihood ratio test and Benjamini-Hochberg correction) between wildtype (n = 128 cells) and *JAK2^V617F^* mutant (n = 93 cells) HSPC1 cells from untreated MF patients (n = 4; excluding PV sample). Only motif accessibility of transcription factors expressed in scRNA-seq data from MPN patients^29^ were considered. Transcription factors involved in biological pathways such as myeloid/erythroid differentiation (pink), NF-κB signaling (teal) and AP-1 complex (JUN/FOS; blue) are highlighted. The horizontal dotted line represents FDR = 0.05, the vertical dotted lines represent absolute Δz-score > 0.25. **e**, Transcription factor footprinting for JUN or NFKB1 motif sites in either wildtype or *JAK2^V617F^* mutant cells within the HSPC1 cluster for untreated or ruxolitinib-treated patients. **f**, Motif accessibility for either JUN or NFKB1 transcription factors for a longitudinal sample (Pt-01) from a patient that progressed from polycythemia vera (PV) to myelofibrosis (MF). Wilcoxon rank sum test. **g**, Volcano plot of the differentially accessible transcription factor motifs (FDR < 0.05 and absolute Δz-score > 0.25; LMM followed by likelihood ratio test and Benjamini-Hochberg correction) between wildtype (n = 312 cells) and mutant (n = 517 cells) HSPC2 cells from untreated MF patients (n = 4; excluding PV sample). Transcription factors involved in biological pathways such as myeloid/erythroid differentiation (pink) and TGF-β signaling (purple) are highlighted. The horizontal dotted line represents FDR = 0.05, the vertical dotted lines represent absolute Δz-score > 0.25. **h**, Correlation between motif accessibility of transcription factors belonging to the STAT family and transcription factors involved in NF-κB signaling within HSPCs (HSPC1 and HSPC2 clusters combined) in untreated MF patients (n = 4; excluding PV sample). Spearman correlation. **i**, Scatter plots illustrating transcription factor motif accessibility correlation in untreated MF HSPCs (HSPC1 and HSPC2) between NFKB1 and STAT1 (left panel) and STAT4 (right panel). Each point represents a single cell, and the lines represent the linear fit for either wildtype (blue) or *JAK2^V617F^* mutant (red). Shadowed areas represent 95% confidence intervals. Linear modeling followed by ANOVA test for comparison between wildtype and mutant slopes, as well as the Rho values representing the goodness of fit for each genotype are shown. **j**, Schematic representation of a *Jak2^RL^* mouse experiment^110^ in which bulk RNA-seq was performed on sorted LSK cells from *Jak2^V617F^* and *Jak2^V617F^*-deleted mice. **k**, Pre-ranked gene set enrichment of differentially expressed genes within the erythroid (FDR = 2.5 x 10^-4^; NES = −1.87; heme metabolism Hallmark gene set) and TNF via NF-κB (FDR = 4.1 x 10^-4^; NES = −1.59) gene sets in *Jak2^V617F^* compared to *Jak2^V617F^-deleted* mouse LSK cells.

Consistently, gene pathway analysis revealed an enrichment in the inflammatory response pathway in mutant HSPC1 cells (Family-wise error rate [FWER] = 0.1031; normalized enrichment score [NES] = 1.375; Hallmark pathway M5932; **Fig. 3b**). In addition to the genes involved in the inflammatory pathway, we observed increased gene accessibility of *S100A12* (**Fig. 3c, left panel,** decreased accessibility of *GTSE1* gene is shown for comparison) encoding a protein involved in pro-inflammatory cytokine upregulation, including TNF through activation of NF-κB signaling^69^. Notably, the pro-inflammatory signature observed in mutant cells was lost upon ruxolitinib treatment (**Extended Data Fig. 5a-b; Extended Data Table 3**), consistent with ruxolitinib resulting in lowered circulating cytokine levels in patients^39,56^.

To further explore the regulatory underpinning of inflammatory phenotypes in HSPCs, we leveraged chromatin accessibility to infer transcription factor activity based on the accessibility of their DNA binding motifs (**see materials and methods**)^70,71^. Comparing wildtype and *JAK2^V617F^*-mutated cells within the HSPC1 cluster, we uncovered a subset of transcription factors that show increased motif accessibility (false discovery rate [FDR] < 0.05 and Δz-score > 0.25) in early mutant HSPCs (**Fig. 3d; Extended Data Table 4**). Transcription factors involved in NF-κB signaling (NFKB1, NFKB2, REL, RELA, and RELB) and in the AP-1 complex (JUN, JUNB, JUND, FOS, FOSB, FOSL1, FOSL2) showed increased accessibility in mutant cells, suggesting that the *JAK2^V617F^* mutation already primes early HSPCs towards a pro-inflammatory cellular state. These findings are consistent with recent studies that have identified NF-κB and STAT3 as essential co-regulatory drivers of inflammation in MPN murine models, resulting in aberrant cytokine signaling from both mutant and non-mutant cells^26,72^. Additionally, NF-κB signaling has been implicated in the regulation of AP-1 transcription factors^73^, and along with STAT3, act as important mediators of an inflammatory regulatory network driving complex transcriptional programs in a variety of human cancers^74,75^. Transcription factor footprinting within the HSPC1 cluster showed increased accessibility surrounding JUN and NFKB1 motifs in *JAK2^V617F^* mutant cells (**Fig. 3e**).

By leveraging longitudinal sampling obtained from an untreated patient with PV that later progressed to MF (Pt-01, see **Extended Data Table 1**), we explored whether the pro-inflammatory phenotype as measured by JUN and NFKB1 motif accessibility preceded the increase in bone marrow fibrosis observed in MF. Indeed, we observed that both JUN and NFKB1 motif accessibilities were increased in *JAK2^V617F^* mutant early HSPCs already at the PV stage (**Fig. 3f**). Importantly, JUN and NFKB1 transcription factor motif accessibility was increased in MF relative to PV (**Fig. 3f**), consistent with an increasingly pro-inflammatory bone marrow environment in the progression from PV to MF. Thus, while pro-inflammatory phenotypes have been linked to extrinsic effects^25,26,72^, and highlighted to affect transcriptional profiles in committed erythroid progenitors^32,76^, our high-resolution mapping of chromatin accessibility in early mutant human HSPCs showed that *JAK2^V617F^* promotes cell-intrinsic pro-inflammatory gene accessibility and transcription factor activity, well before commitment towards the erythroid fate. myeloid/erythroid differentiation (LYL1^77,78^, SNAI1^79^, TCF4^80^, and MESP1^81^) also exhibited increased accessibility of their DNA-binding motifs in mutant cells (**Fig. 3d**), suggesting that epigenetic priming towards a myeloid cell fate is present at early stages of *JAK2^V617F^*-mutated hematopoiesis. We also observed modestly decreased motif accessibility for a group of transcription factors involved in stem cell quiescence (NFYA/B/C^82,83^, YY1^84^, and SP2^85,86^) in mutant cells (**Fig. 3d**). The decrease in potential activity of these transcription factors may underlie the increased hematopoietic output of mutant cells relative to their wildtype counterparts, as suggested by increased mutant fractions in more differentiated progenitor clusters in the myeloid lineage.

To uncover further changes in transcription factor activity within later HSPCs, we performed differential transcription factor motif accessibility between wildtype and mutant cells within the HSPC2 cluster (**Fig. 3g; Extended Data Table 4**). Consistent with our observations in the earlier HSPC1 cluster, differential transcription factor motif accessibility highlighted increased activity of transcription factors associated with myeloid/erythroid differentiation (e.g., ID3^87^, MESP1^81^, SNAI1^79^, LMO2^88–91^, TCF4^80^, ZEB1^92^, and LYL1^77,78^) in mutant cells, suggesting that the erythroid bias is strengthened in later HSPCs, likely underlying erythroid priming^6^. Notably, LMO2, a key factor in erythroid differentiation^88–91^, has been postulated as a direct non-signaling target of JAK2 through nuclear translocation and histone modulation^48^, thus serving as a potential direct link between the mutated genotype and erythroid-biased HSPC priming. A second group of transcription factors also associated with myeloid and erythroid differentiation showed a more modest yet significant increase in *JAK2^V617F^* mutant HSPCs (e.g. TAL1^78,93^, RUNX1^94,95^, TFCP2^96^, NFIA^97^, NFIB^98^, and CBFB^99^; **Fig. 3g**). Of note, RUNX1 is known to drive the initial commitment towards myeloid differentiation in hematopoiesis^9,95,100^ and may underlie the myeloid priming observed in myelofibrosis.

Consistent with mutant cells in the HSPC1 cluster, HSPC2 mutant cells also exhibited a decrease in the accessibility of transcription factor motifs implicated in stem cell quiescence (NFYA/B/C^82,83^, YY1^84^, YY2^101^, GATA2^102–104^ and HLF^105^; **Fig. 3g**). Additionally, HSPC2 mutant cells showed an increase in accessibility of transcription factor motifs within the TGF-β signaling pathway (TGIF2, SMAD1, SMAD4, and SMAD9), which is thought to play an important role in the development of marrow fibrosis^106–109^. Of note, ruxolitinib treatment abrogated the cell-intrinsic differences observed in transcription factor motif accessibility between wildtype and *JAK2^V617F^-mutated* HSPCs (**Extended Data Fig. 5c-d; Extended Data Table 4**), demonstrating that the changes observed in untreated patients are mediated by the *JAK2^V617F^* constitutive activation.

To link the activation of NF-κB signaling to canonical JAK2 downstream targets, we correlated NF-κB-related transcription factor motif accessibility with members of the STAT family transcription factor motifs in HSPCs. We found increased correlation of STAT1, STAT3, STAT5A, and STAT5B motif accessibility with NF-κB factor motif accessibility, consistent with JAK2 activation of canonical STAT targets (**Fig. 3h; Extended Data Fig. 5e**). In contrast, non-canonical JAK2 targets STAT2, STAT4, and STAT6 showed reduced correlation values with NFKB1 compared to canonical JAK2 target STATs (**Fig. 3h; Extended Data Fig. 5e**), suggesting that JAK2-mediated activation underlies the increased activity of the NF-κB-associated transcription factors. Indeed, although both wildtype and *JAK2^V617F^* mutant HSPCs show a positive correlation between STAT1 and NFKB1 motif accessibility, mutant cells display a higher correlation coefficient (**Fig. 3i, left panel**; *P* = 7.1 x 10^-4^; Spearman correlation), while no differences were observed in the correlation between STAT4 and NFKB1 (**Fig. 3i, right panel**; *P* = 0.21; Spearman correlation). Thus, activation of STAT1, STAT3, STAT5A and STAT5B downstream of JAK2^V617F^ might drive the observed NF-κB signal.

To validate the inflammatory and myeloid/erythroid signatures observed in mutant HSPCs, we leveraged bulk RNA-seq data of LinM^-^, Sca-1^+^, c-Kit^+^ (LSK) progenitor cells from a novel Dre-*rox*, Cre-*lox* dual recombinase *Jak2^V617F^* mouse model that allows for sequential knock-in followed by knock-out of the mutated allele^110^ (*Jak2^Rox/Lox^/Jak2^RL^*; **Fig. 3j**). Consistent with our findings in human patient samples showing erythroid priming and a cell-intrinsic pro-inflammatory signals, gene set enrichment analysis revealed an enrichment of the heme metabolism gene set which includes key erythroid transcription factors (FDR = 2.5 x 10^-4^; NES = −1.87), and the tumor necrosis factor (TNF) signaling via NF-κB gene set (FDR = 4.1 x 10^-4^; NES = −1.59) in *Jαk2^V617F^* compared to *Jαk2^V617F^-deleted* murine LSK cells (**Fig. 3k**), supporting the causality of the *JAK2^V617F^* mutation in driving the observed phenotypes.

Collectively, gene accessibility and transcription factor motif accessibility comparisons revealed a subset of early HSPCs displaying cell-intrinsic pro-inflammatory phenotypes, as well as erythroid lineage priming in *JAK2^V617F^* mutant versus wildtype human HSPCs.

### *JAK2^V617F^* results in cell type-specific epigenetic changes in committed erythroid and megakaryocytic progenitors

We next sought to define the epigenetic changes in the erythroid and megakaryocytic progenitors, the cell types undergoing significant clonal expansion of *JAK2^V617F^-mutated* cells (**Fig. 2g**). When assessing transcription factor motif accessibility changes in the EP1 cluster, we found increased STAT5A and STAT5B transcription factor motifs (**Fig. 4a; Extended Data Table 4**), with increased accessibility of STAT1, STAT3, STAT5A, and STAT5B motifs in the late erythroid progenitor (EP2) cluster (**Extended Data Fig. 6a; Extended Data Table 4**). These results highlight the cell type-specific degree of activation of canonical downstream targets of JAK2, varying between HSPCs and between early and late EPs, which might drive the increased fitness of mutated erythroid progenitors^111,112^. In addition, we observed increased accessibility of multiple myeloid/erythroid transcription factor binding motifs (TCF4^80^, ID3^87^, MESP1^81^, TFCP2^96^, POU2F1^113^, THRB^114^, THRA^115^, ZEB1^92^, LMO2^88–91^, and DDIT3^116^; **Fig. 4a**), consistent with our previous observations in early HSPCs, pointing to a *JAK2^V617F^* effect that remains consistent across erythroid differentiation. In contrast, PU.1 (*SPI1*), a critical regulator of cell fate decisions between myeloid and lymphoid lineages^117^, exhibited decreased motif accessibility in mutant cells. Recent work has shown that downregulation of PU.1 induces gene expression changes that promote myeloid-biased stem cells and erythroid differentiation^118,119^. Thus, increased erythroid-associated and STAT transcription factor activity may result in increased fitness underlying the expansion of mutated erythroid progenitors.

**Fig. 4.**
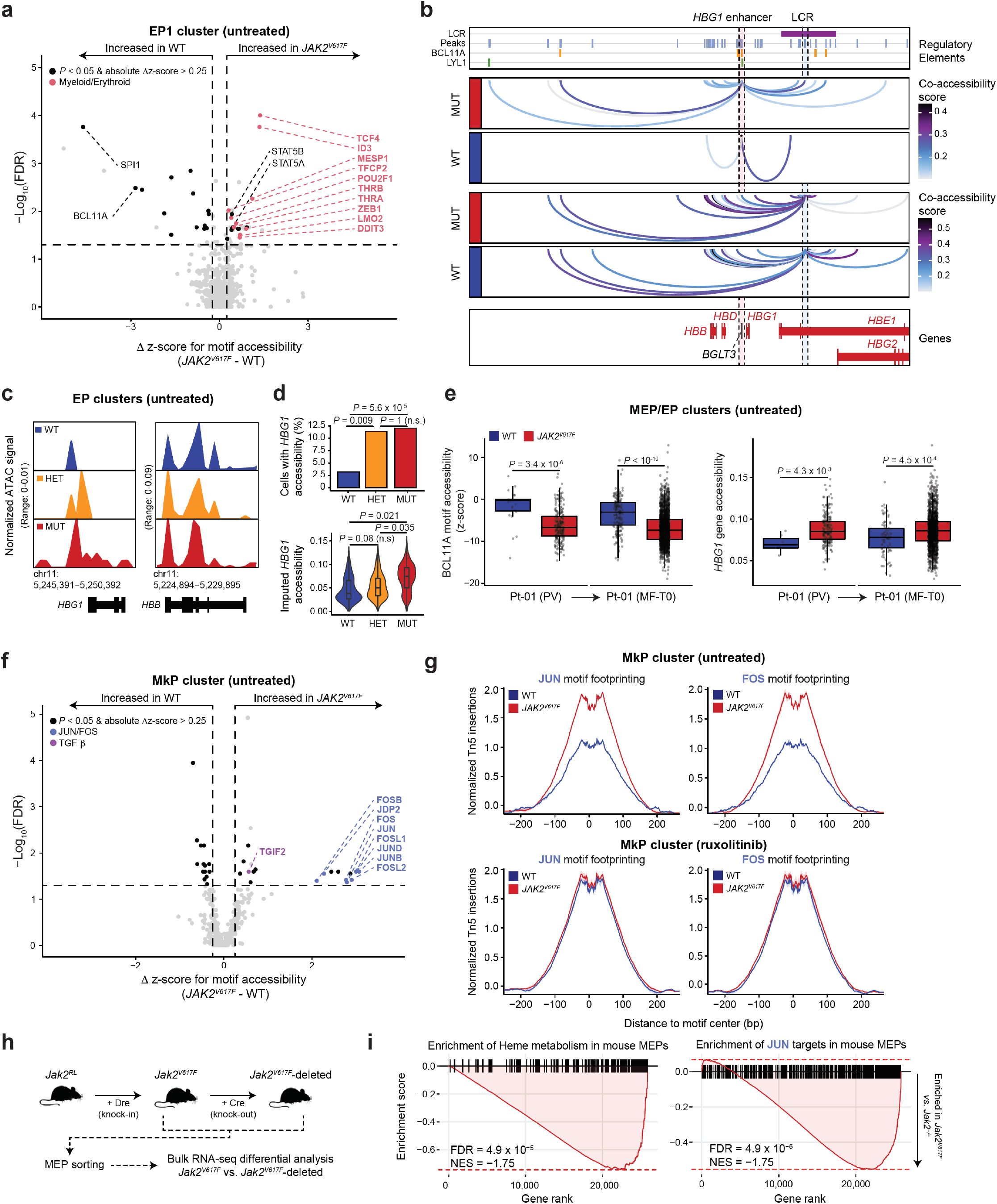
GoT-ChA reveals *JAK2^V617F^*-driven epigenetic dysregulation of hemoglobin locus in EPs and increased JUN/FOS signaling in MkPs. **a**, Differential transcription factor motif accessibility between wildtype (n = 121) and mutant (n = 703) cells within the EP1 cluster of untreated MF patient samples (n = 4). Horizontal dotted line represents FDR = 0.05; vertical dotted lines represent absolute Δz-score > 0.25; transcription factors involved in erythroid differentiation are highlighted in pink. Linear mixture model (LMM) followed by likelihood ratio test and Benjamini-Hochberg correction. **b**, Examples of co-accessibility measurements (correlation > 0.1 and FDR < 0.05; Wilcoxon rank sum test followed by Benjamini-Hochberg correction) of regulatory elements surrounding the *HBG1* locus for wildtype (n = 121 cells; blue) and mutant (n = 703 cells; red) cells in the EP1 cluster. Peaks (light blue), the hemoglobin locus control region (LCR, purple), and transcription factor motifs for BCL11A (red) and LYL1 (yellow) are indicated. Co-accessibility measurements centered in two peaks of interest are indicated via shaded dotted lines for an enhancer controlling *HBG1* expression (shaded in red) on top, and a comparison peak within the LCR (shaded in blue). Gene location and shaded regions indicating location of highlighted peaks are shown at bottom. **c**, Chromatin accessibility coverage tracks comparing accessibility between homozygous wildtype (blue), heterozygous (yellow), and homozygous mutant (red) cells within the EP1 cluster for the *HBG1* and *HBB* genes. **d**, Quantification of the proportion of cells showing accessibility of the *HBG1* gene (upper panel; Fisher exact test) or the imputed gene accessibility value for either homozygous wildtype, heterozygous or homozygous mutant cells (bottom panel; LMM followed by likelihood ratio test). **e**, BCL11A motif accessibility and *HBG1* gene accessibility scores for longitudinal sample (Pt-01) from a patient progressing from polycythemia vera (PV) to myelofibrosis (MF). Wilcoxon rank sum test. **f**, Differential transcription factor motif accessibility between wildtype (n = 95 cells) and mutant (n = 542 cells) within the MkP cluster of untreated MF patients (n = 4). A subset of TFs that exhibited enrichment in mutant cells are highlighted by pathway involvement: JUN/FOS signaling in blue, TGF-β in purple. Horizontal dotted line represents FDR = 0.05; vertical dotted lines represent absolute Δz-score > 0.25. LMM followed by likelihood ratio test and Benjamini-Hochberg correction. **g**, Transcription factor footprinting for JUN and FOS comparing wildtype (blue) and mutant (red) in untreated (n = 4; top) and treated (n = 3; bottom) MF patient samples. **h**, Schematic representation of a *Jak2^RL^* mouse experiment^110^ in which bulk RNA-seq was performed on sorted MEP cells from *Jak2^V617F^* and *Jak2^V617F^-deleted* mice. **i**, Gene set enrichment analysis illustrating a depletion of JUN targets and heme metabolism pathway (Hallmark M5945) in *Jak2^V617F^-deleted* compared to *JAK2^V617F^* mouse MEPs (NES = normalized enrichment score).

Of note, BCL11A was one of the top transcription factors noted to have decreased motif accessibility in mutated erythroid progenitors (FDR = 0.0032, Δz-score = −2.84; **Fig. 4a**). BCL11A is required for repression of the γ-globin gene (*HBG1*)^120^, and loss of BCL11A is sufficient for increased expression of *HBG1*^121,122^, a component of fetal hemoglobin (HbF). To further explore the changes in the regulatory elements controlling the hemoglobin locus, scATAC-seq data is particularly well suited for defining pairs of co-accessible regions within the same single cell, which reflect coordinated activation of regulatory elements^123^. By utilizing single-cell genotyping with GoT-ChA, we performed co-accessibility analysis of the hemoglobin locus region in either wildtype or *JAK2^V617F^* mutant erythroid progenitors. We found that an enhancer region (ENCODE accession EH38E1516933) upstream of *HBG1* showed increased co-accessibility in *JAK2^V617F^* mutant erythroid progenitors when compared to their wildtype counterparts (**Fig. 4b, red highlighted region**), while another enhancer region located within the locus control region^124^ controlling the β-globin expression showed consistent co-accessibility across wildtype and *JAK2^V617F^* mutant cells (**Fig. 4b**, **grey highlighted region**). These data provide further insight into reports that have shown an increase in HbF in MPN patients^125–127^. By leveraging GoT-ChA and comparing genotyped cells, we observed a consistent increase in the accessibility of *HBG1* (**Fig. 4c**) proportional to the *JAK2^V617F^* allelic burden, while no apparent changes were observed in accessibility of the β-globin gene (*HBB*; **Fig. 4c**). Indeed, we observed an increase in the proportion of mutant cells in the EP1 cluster showing accessibility of the *HBG1* gene (**Fig. 4d, upper panel**; Fisher exact test), while *HBG1* gene accessibility increased proportionally to the mutated allele burden (**Fig. 4d, bottom panel**; LMM followed by likelihood ratio test). By leveraging longitudinal sampling of Pt-01 (PV → MF), we observed that decreased BCL11A motif accessibility as well as increased *HBG1* gene accessibility are already present at the PV stage (**Fig. 4e**; Wilcoxon rank sum test) and remain upon progression to MF. While increased HbF levels often result from defective erythropoiesis, our data reveals that cell-intrinsic regulatory changes in *JAK2^V617F^* mutant erythroid progenitors may also promote HbF and precede the onset of marrow fibrosis. In further support of a cell-intrinsic mechanism, these changes were reverted by ruxolitinib treatment (**Extended Data Fig. 6b-d**), indicating the requirement of constitutive JAK2^V617F^ activity for the increased accessibility of *HBG1* and decreased accessibility of BCL11A binding motifs.

A key clinical feature of myelofibrosis is a dramatic increase in megakaryocytes, which are thought to be one of the main cellular drivers of marrow fibrosis via pro-fibrotic cytokine and growth factor signaling^109,128–130^. Differential transcription factor motif accessibility in MkPs revealed an increased activity of JUN and FOS family proteins (**Fig. 4f; Extended Data Table 4**), which have recently been implicated in inflammation^74^ and as important mediators of fibrosis in various conditions, including myelofibrosis^43,44^. In contrast, differential transcription factor motif accessibility signals were reverted by ruxolitinib treatment (**Extended Data Fig. 6e; Extended Data Table 4**), consistent with mutated versus wildtype motif footprinting analysis for JUN (**Fig. 4g**). To validate the dependency of JUN/FOS activation on *JAK2^V617F^*, we again interrogated the reversible *Jak2^RL^* mouse model^110^ (**Fig. 4h**). Consistent with our findings in human myelofibrosis, differential bulk RNA-seq analysis of sorted megakaryocytic-erythroid progenitors (MEPs) showed depletion of an erythroid gene set as well as JUN targets upon reversal of the *Jak2^V617F^* allele (**Fig. 4i**), demonstrating that *Jak2^V617F^* is required for increased JUN activity in MEPs. Taken together, these data demonstrate cell type-specific and cell-intrinsic alterations in the epigenetic landscape leading to aberrant erythropoiesis and increased pro-fibrotic JUN/FOS activation in megakaryocyte progenitors in MF.

### GoT-ChA with select antigen profiling allows mitochondrial-based genotyping imputation and protein measurement integration

To expand the reach of GoT-ChA for multi-modality single-cell sequencing, we integrated it with ASAP-seq^38^, a method that assays genome-wide chromatin accessibility simultaneously with targeted protein expression utilizing fixed whole cells (rather than nuclei in standard scATAC-seq). We applied the combined method to two MF patient samples, Pt-02 (untreated) and Pt-06 (ruxolitinib treated). Genotyping of the *JAK2^V617^* locus was available for 2,663 out of 11,457 (23.2%) and for 489 out of 2,928 (16.7%) cells for Pt-02 and Pt-06, respectively (**Extended Data Fig. 7a**), while retaining chromatin accessibility quality (**Extended Data Fig. 7b-c**). We note a decrease in genotyping rate with GoT-ChA-ASAP (23.2% and 16.7% for Pt-02 and Pt-06, respectively) compared to when the samples were profiled with GoT-ChA alone (51.8% and 27.8% for Pt-02 and Pt-06, respectively), likely due to the fixation conditions required for ASAP-seq. However, by assaying entire cells rather than isolating nuclei during sample preparation, ASAP-seq results in a massively increased coverage of the mitochondrial genome (**Fig. 5a**)^38^. We hypothesized that *JAK2^V617F^* mutant cells might harbor mitochondrial variants, as previously observed^31^ (**Extended Data Fig. 7d**), that could be utilized for imputation of genotypes in cells where the *JAK2* locus was not captured directly. Of the two GoT-ChA-ASAP samples, only Pt-02 showed two mitochondrial variants (12,786 A>G and 3,834 G>A) at high heteroplasmy in *JAK2^V617F^* cells (average heteroplasmy of 77.7% and 18.1%, respectively), while exhibiting low heteroplasmy in wildtype cells (average heteroplasmy of 7.85% and 1.47%, respectively; **Fig. 5b**). While both mitochondrial variants were in phase with the *JAK2^V617F^* mutation, they were mutually exclusive, suggesting that they were acquired early after the *JAK2^V617F^* mutation.

**Fig. 5.**
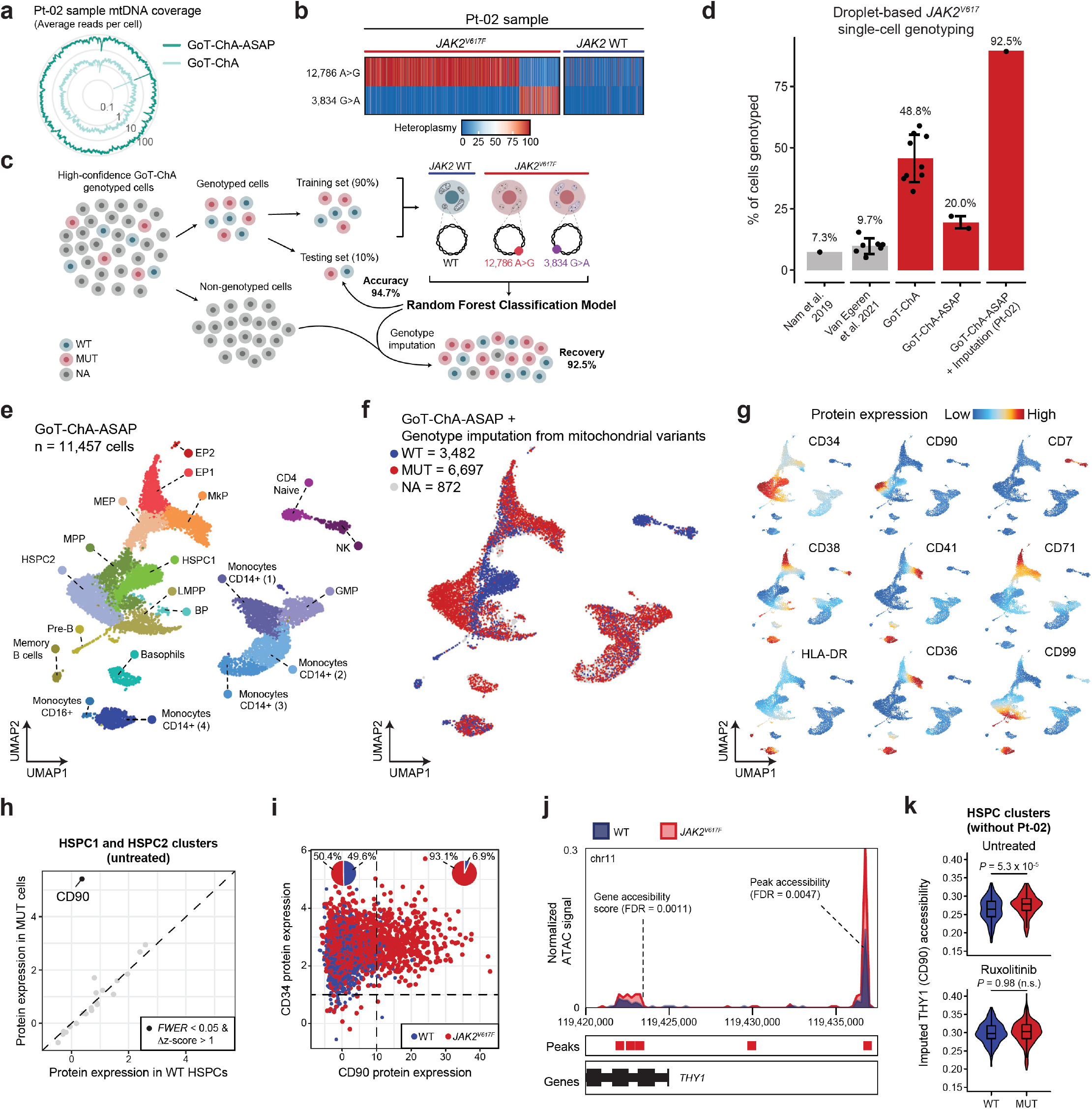
GoT-ChA integration with ASAP-seq enables mitochondrially informed genotype imputation and surface protein measurements in primary human samples. **a**, Depiction of the overall mitochondrial genome coverage from sample Pt-02 processed with GoT-ChA-ASAP compared to the same sample with the standard GoT-ChA protocol. The average number of reads per cell covering each region is shown. **b**, Heatmap of heteroplasmy of mitochondrial variants per cell for patient sample Pt-02, illustrating the presence of two variants (12,786 A>G and 3,834 G>A) that segregate in phase with GoT-ChA genotyping. **c**, Schematic representation detailing the process of genotype imputation using a random forest classifier. GoT-ChA genotyped cells were separated into a training set (90% of cells, downsampled for even numbers of mutant and wildtype cells) and a test set (10% of cells). The classifier resulted in an imputed genotyping accuracy of 94.7% on the test set and, when applied to non-genotyped cells in the same sample, increased the effective rate of genotyping in the sample from 23.2% to 92.5%. **d**, Percent of *JAK2^V617^* genotyped cells across multiple studies and protocols utilizing high-throughput droplet-based technologies. Bars represent the mean; each dot represents a sample and error bars represent the standard deviation. **e**, Uniform manifold approximation and projection (UMAP) of cells from sample Pt-02 processed with GoT-ChA-ASAP, illustrating the expected stem cell and committed progenitor populations in early hematopoiesis (n = 11,457 cells). HSPC = hematopoietic stem and progenitor cells; MPP = multi-potent progenitors; GMP = granulocyte monocyte progenitors; MEP = megakaryocyte-erythrocyte progenitors; EP = erythroid progenitors; MkP = megakaryocytic progenitors; LMPP = lymphoid-myeloid pluripotent progenitors; Pre-B = precursor B cells; NK = natural killer cells; BP = basophil progenitors. **f**, UMAP as seen in **e**, colored by GoT-ChA genotyping of the *JAK2^V617^* locus after mitochondrial-based imputation: wildtype (WT; n = 3,482 cells; blue), mutant (MUT; n = 6,697 cells; red), and not assignable (NA; n = 872 cells; grey). **g**, UMAP as seen in **e**, colored by protein expression of cell surface markers, illustrating successful measurement of protein expression with GoT-ChA-ASAP. **h**, Differential protein expression between wildtype and mutant cells within HSPC1 and HSPC2 clusters. Wilcoxon rank sum test followed by Bonferroni correction. Black dots represent differentially expressed (family wise error rate [FWER] < 0.05 and absolute Δz-score > 1) cell surface proteins. **i**, Scatter plot comparing protein expression of CD34 and CD90. Inset pie charts show the enrichment of *JAK2^V617F^* mutant cells in each CD34^+^quadrant. **j**, Chromatin accessibility coverage track of the CD90 gene (*THY1*) and a proximal upstream regulatory element is shown for either wildtype (blue) and mutant cells (red) from the HSPC1 and HSPC2 clusters. Wilcoxon rank sum test followed by Benjamini-Hochberg correction. **k**, CD90 (*THY1* gene) chromatin accessibility scores of HSPCs (HSPC1 and HSPC2 clusters) for untreated MF samples (upper panel; *P* = 5.3 x 10^-5^; linear mixture model (LMM) modeling patient identity as random effects, followed by likelihood ratio test) excluding Pt-01 (PV) and sample Pt-02, or ruxolitinib-treated samples (P = 0.98; LMM modeling patient identity as random effects, followed by likelihood ratio test).

To leverage the clonal phasing of mitochondrial variants and *JAK2^V617F^*, we developed a random forest classifier to impute missing *JAK2* genotypes based on heteroplasmy levels. We trained the classifier on a random sampling of 90% of Pt-02 genotyped cells (training set) and assessed performance with the remaining 10% of genotyped cells (test set), resulting in a genotyping accuracy of 94.7% (**Fig. 5c**). Implementation of the classifier on non-genotyped cells resulted in a striking increase in genotyped cells from 23.2% to 92.53% of cells (**Fig. 5d; Extended Data Fig. 7e**). Thus, while standalone GoT-ChA already demonstrated a significant improvement for genotyping loci that were previously challenging due to low expression and mutation location, the additional implementation of an imputation classifier built on phased mitochondrial variants yielded the ability to accurately genotype nearly all cells within the sample.

Cell clustering of the Pt-02 sample based on chromatin accessibility alone, and therefore agnostic to genotyping, protein, or mitochondrial data, resulted in the expected cell clusters (**Fig. 5e**) identified via accessibility of canonical marker genes (**Extended Data Fig. 7f-g**). Importantly, while mutant and wildtype cells intermingled in most progenitor subtypes, HSPC clusters 1 and 2 were predominantly enriched by wildtype and mutant cells, respectively, suggesting a discrete enrichment of later HSPCs (HSPC2) with *JAK2^V617F^-mutated* cells in this sample (**Fig. 5f**).

We further leveraged GoT-ChA-ASAP for simultaneous measurement of protein expression, applying a panel of 21 cell surface protein markers to orthogonally validate cluster assignments (**see materials and methods**). Progenitor clusters HSPC1, HSPC2, and MPP showed increased CD34 and decreased CD38 staining, while MkPs showed increased CD41 and CD36, EPs showed high CD71, lymphoid clusters showed high CD99 staining, and T-cells showed high CD7 levels (**Fig. 5g**). Differential protein expression comparisons showed that mutant HSPCs exhibited higher expression of CD90 than their wildtype counterparts (**Fig. 5h**; Δz-score > 1 and FWER < 0.05; Wilcoxon rank sum test followed by Bonferroni correction), while Pt-06, undergoing ruxolitinib treatment at the time of sample collection, showed no significant changes in CD90 protein expression levels between wildtype and *JAK2^V617F^* cells within HSPCs (**Extended Data Fig. 7h**). Indeed, Pt-02 CD34^high^, CD90^high^ HSPC cells were comprised of 93.1% mutant cells compared to 50.4% of CD34^high^, CD90^low^ cells (**Fig. 5i**). Concordantly, Pt-02 mutant HSPCs exhibited an increase in gene accessibility of the CD90 gene (*THY1;* FDR = 1.1 x 10^-3^), as well as in accessibility of a proximal upstream regulatory element (FDR = 4.7 x 10^-3^) relative to wildtype cells (**Fig. 5j**). Importantly, we observed increased accessibility of the CD90 gene in HSPCs in the other untreated MF samples in this cohort (**Fig. 5k, upper panel**; *P* = 5.3 x 10^-5^; LMM followed by likelihood ratio test, HSPC1 and HSPC2 combined; see **Extended Data Fig. 7i** for per patient analysis) while ruxolitinib treatment reverted this effect (**Fig. 5k, bottom panel**; *P* = 0.98; LMM followed by likelihood ratio test, HSPC1 and HSPC2 combined). CD90 is a cell surface marker expressed in a subset of primitive HSPCs^131^. Indeed, expansion of the CD90^high^ HSPC cell population has been reported in PV^23^. Furthermore, Rodriguez-Meira and colleagues also noted aberrant expression of CD90 in homozygous *JAK2^V617F^* MPN HSPCs using TARGET-seq^25^. However, due to the lower throughput of plate-based methods, this observation was supported by only five cells, demonstrating the advantage of high-throughput single-cell multi-omics integration.

Overall, these results demonstrate the capability of GoT-ChA to deliver a highly multi-modal single-cell platform to link mutated genotypes with mitochondrial variants and cell surface proteins, together with chromatin accessibility, enabling discovery of clonal changes across multiple layers of information in a single, high-throughput, unified assay.

## DISCUSSION

A critical challenge in studying the phenotypes of clonal expansions, in both healthy tissues and overt malignancies, is that primary human samples are often composed of an admixture of wildtype and mutant cells. Thus, precision mapping of genotypes to phenotypes is obscured. Additionally, bulk population measurements likely aggregate heterogeneous groups of cells, hindering the identification of cell type- or cell state-specific phenotypes arising from the presence of somatic mutations. To circumvent these limitations, single cell simultaneous capture of genotypes together with phenotypic information is required, enabling intra-sample, cell type-specific mutant to wildtype comparisons. Application of such single cell multi-omic approaches have shown that somatic mutations in human tissues exert a differing phenotypic effect as a function of cell state^5,6,24,25,28–33^.

While droplet-based single cell technologies allow high-throughput linkage of cDNA-captured mutated genotypes with phenotypes^29,31^, similar high-throughput methods for linking somatic genotypes with epigenetic profiles were lacking. Furthermore, cDNA-based capture results in limiting dependencies on target gene expression and the distance of the locus from transcript end. For example, these dependencies led to inefficient capture of the lowly expressed *JAK2^V617F^* locus (<10% of cells^29,31^), requiring gDNA capture via lower throughput plate-based single cell sequencing^25,33,132^. The dependency on gene expression also limits the application to archival frozen tissues, as nuclei isolation further decreases the number of available mRNA molecules for genotyping. Collectively, these limitations restrict the ability to profile key lowly expressed mutations, as well as mutations leading to nonsense-mediated mRNA decay or affecting non-coding regions.

GoT-ChA addresses this challenge by delivering droplet-based, broadly available, high-throughput joint capture of genotypes and chromatin accessibility. We further show that GoT-ChA can be readily integrated with protein and mitochondrial DNA capture, enabling robust linkage of somatic genotypes to a variety of signals at single cell resolution. As GoT-ChA is based on gDNA rather than cDNA capture, it also obviates the limiting dependencies on mutated locus expression and location. Thus, GoT-ChA enables the interrogation of somatic mutations throughout the genome, and radically expands the range of human biological phenomena that can be investigated for epigenetic deregulation due to somatic mutations. Importantly, the ability to apply GoT-ChA to nuclei opens the possibility for application to archived frozen solid tissues or tumors, critical for the exciting emerging field of clonal mosaicism across human tissues^133–142^.

To leverage the unique ability to chart the impact of somatic mutation on epigenetic differentiation landscapes, we focused on *JAK2^V617F^-driven* clonal expansions in primary human samples from PV and MF patients. These data revealed that the epigenetic consequences of the *JAK2^V617F^* mutation are highly cell state dependent. Indeed, the frequency of mutated cells expanded at the stage of committed erythroid and megakaryocytic progenitors, consistent with clinical phenotypes^39^, and demonstrates that the clonal representation of this mutation varies by differentiation stage and fate. Notably, the ability to profile single-cell chromatin landscapes allowed to examine patterns of HSPC priming and demonstrated that increases in the frequencies of mutated committed progenitors were heralded by increased accessibility of erythroid transcription factor motifs already in mutated HSPCs, consistent with aberrant lineage priming in MPN initiating cells.

*JAK2^V617F^* myeloproliferation is characterized by the presence of an inflammatory microenvironment, driving bone marrow fibrosis and extramedullary hematopoiesis^7,20,21^. Previous work has linked JAK2-mediated activation of STAT1 and STAT3 to increased NF-κB signaling in mouse models^26,72^, and highlighted cell-extrinsic effects of the microenvironment. Here, we show that the *JAK2^V617F^* mutant HSPCs also display epigenetic profiles that are consistent with cell-intrinsic pro-inflammatory phenotypes, with increased motif accessibility of NF-κB-, AP-1-, and TGF-β-associated transcription factors in mutant cells. The observation that pro-inflammatory phenotypes are, at least to some degree, linked directly to *JAK2^V617F^* in a cell-intrinsic fashion opens an avenue for potential combined therapeutic strategies for mutant-specific targeting, aimed at both *JAK2^V617F^* constitutive activation as well as pro-inflammatory signaling in mutant HSPCs. In another striking demonstration of cell-type specificity in mutational impact, *JAK2^V617F^* megakaryocytic progenitors, which produce mature megakaryocytes thought to drive characteristic marrow fibrosis through pro-fibrotic cytokine signaling^109,128–130^, showed a pro-inflammatory landscape specific for AP-1 transcription factor activity, which has been linked with the fibrotic clinical phenotype of MF. In contrast, early HSPCs (HSPC1) showed broad, pro-inflammatory signature characterized not only by increased AP-1-, but also NF-κB- and TGF-β-associated transcription factor motif accessibility. The study of paired samples from a PV patient who progressed to MF showed that many of the changes described above, including increased pro-inflammatory NF-κB and AP-1 motif accessibility in HSPCs and aberrant regulation of the γ-globin locus in EPs, are already evident long before significant marrow fibrosis occurs, providing human data support to the causal role of JUN/FOS in inducing fibrosis. These data suggest that *JAK2^V617F^*-mediated inflammation and fibrosis results from a complex interplay between cell-extrinsic^25,26,72^ and cell-intrinsic effects that vary across different progenitor populations.

Interestingly, we observed a near complete loss of differential accessibility signals between mutant and wildtype cells within patients treated with ruxolitinib therapy, including reversion of intrinsic pro-inflammatory phenotypes and differentiation biases. However, ruxolitinib treatment does not eliminate *JAK2^V617F^* mutant HSPCs. This is in line with the observation that while ruxolitinib reduces splenomegaly and disease symptoms resulting in overall improvement in quality of life^39,143^, it fails to prevent disease progression or eliminate the mutated clones in MPNs^40–42^. Thus, while ruxolitinib treatment appears to abrogate *JAK2^V617F^*-mediated shifts in the epigenetic landscape of HSPCs, mutated cells may continue to promote disease progression through clonal evolution, even in the context of JAK inhibition^144–147^. Thus, improved JAK2 inhibition for elimination of mutated cells may be critical for the prevention of disease progression^110^.

We note key limitations to the GoT-ChA framework. While our cell mixing study directly testing heterozygous genotyping demonstrated a 34% rate of complete allelic capture, incomplete capture of targeted alleles during in-droplet genotyping reduces the capacity for classification of heterozygous cells. While MF patients tend to have homozygous *JAK2^V617F^* mutations^148–153^, allelic dropout will result in misclassification of heterozygous cells as either homozygous wildtype or mutant. Nonetheless, genotype misclassifications would dilute the strength of the biological differences between genotypes, and thus the differential epigenomic alterations reported here likely serve as a lower bound to effect sizes. Further, application of GoT-ChA-ASAP in samples with in-phase mitochondrial variants may allow for inference genotyping, decreasing the impact of allelic dropout. In addition, although our multi-modal approach expanding GoT-ChA allows to link genotype to both epigenetic changes and to protein expression levels, it does not provide simultaneous capture of transcriptional information, thus limiting the assessment of mutational impact on gene expression. While the 10x Genomics Multiome platform obtains both chromatin accessibility and gene expression information, the cell barcoding reaction does not utilize in-droplet PCR and thus precludes the usage of GoT-ChA. However, development of alternative multiomic technologies that retain an in-droplet barcoding PCR, such as ISSAAC-seq^154^ that utilizes the 10x Genomics scATAC-seq platform as a foundation for simultaneous scRNA-seq, provides promising avenues for linking gene expression changes to mutation-specific epigenetic alterations assayed via GoT-ChA. Furthermore, given that the transposase-enabled scATAC-seq has been leveraged to expand the range of assayable epigenomic profiles to other modalities such as histone modifications^155,156^, GoT-ChA has the potential to link genotypes with additional epigenetic features and therefore provide a more comprehensive understanding of the cellular phenotypes driven by somatic mutations.

Collectively, we report a powerful novel single-cell multiomic approach that allows for the direct investigation of the impact of somatic mutations on chromatin accessibility in primary human patient samples. These data show that the *JAK2^V617F^* somatic mutation, central to MPN pathogenesis, leads to epigenetic rewiring, in a cell-intrinsic and cell type-specific manner. These results, thus, demonstrate the power of joint single-cell capture of genotypes and epigenomes for a high-resolution study of clonal outgrowths in primary human tissue. We propose that GoT-ChA may be of particular importance to the emerging field of clonal mosaicism, now understood to be ubiquitous across the human body^133,134,140^. In non-malignant clonal mosaicism, previous investigations have been largely limited to genotyping, due to the inability to separate admixtures of wildtype and mutant cells for genotype-phenotype inferences in primary human samples. We envision that GoT-ChA will thus serve as a foundation for broad future explorations to uncover the critical link between mutated somatic genotypes and epigenetic alterations across human clonal outgrowths in malignant and non-malignant contexts.

## ACKNOWLEDGMENTS

R.M.M. is supported by a Medical Scientist Training Program grant from the National Institute of General Medical Sciences of the National Institutes of Health under award number T32GM007739 to the Weill Cornell/Rockefeller/Sloan Kettering Tri-Institutional MD-PhD Program and by the Weill Cornell Medicine NYSTEM Training Program under award number C32558GG. F.I. is supported by the American Society of Hematology Fellow-to-Faculty Scholar Award. A.J.D. is a William Raveis Charitable Fund Physician-Scientist of the Damon Runyon Cancer Research Foundation (PST-24-19) and has received funding from the American Association of Cancer Research and the American Association of Clinical Oncology. A subset of biospecimens and data for this work were provided through the Hematological Malignancies Tissue Bank, which is administered and functions under the auspices of the NCI-designated Tisch Cancer Institute at the Icahn School of Medicine at Mount Sinai. R.C. is supported by Lymphoma Research Foundation and Marie Skłodowska-Curie fellowships. R.L.L. is supported by a Leukemia & Lymphoma Society Specialized Center of Research grant and a National Cancer Institute award (P01 CA108671). D.A.L. is supported by the Burroughs Wellcome Fund Career Award for Medical Scientists, Valle Scholar Award, Leukemia Lymphoma Scholar Award, Mark Foundation Emerging Leader Award, and the National Institutes of Health Director’s New Innovator Award (DP2-CA239065). This work was also supported by the Tri-Institutional Stem Cell Initiative, the National Heart Lung and Blood Institute (R01HL145283; R01HL157387-01A1) and the National Human Genome Research Institute, Center of Excellence in Genomic Science (RM1HG011014). This work was enabled by the Weill Cornell Flow Cytometry Core. We thank Dr. Ari Melnick (Weill Cornell Medicine) for a critical review of the manuscript.

## AUTHOR CONTRIBUTIONS

R.M.M., F.I., and D.A.L. conceived the project, devised the research strategy, and analyzed the data. R.M.M., F.I., E. P.M., R.C., P.S., and D.A.L. developed GoT-ChA. R.M.M., F. I., S.K., and D.A.L. developed the analytical pipelines for processing GoT-ChA data. M.S., S.E.G-B., J.A., R.H., B.M., and O.A.-W. conducted a database search and retrieved patient samples for experimental use. R.M.M., S.G., and L. M. performed the experiments. R.M.M., F.I., S.K., T.P., and R.R performed the computational analyses. A.D. and R.L.B. performed the *in vivo* murine bulk RNA-seq experiments. R.M.M., F.I., and D.A.L. wrote the manuscript. R.M.M., F.I., S.K., T.P., A.D., R.L.B., E.P.M., M.S., R.R., S.G., L.M., R.H., R.C., O.A.-W., P.S., B.M., R.L.L., and D.A.L. helped to interpret results. R.M.M., F.I., and D.A.L. acquired funding for this work. All authors reviewed and approved the final manuscript.

## CODE AVAILABILITY

The code used for raw data processing and noise correction approaches for the genotyping data obtained through GoT-ChA, as well as functions for downstream differential gene accessibility, transcription factor motif accessibility and plotting of co-accessibility will be made available as part of the Gotcha R package upon publication.

## DATA AVAILABILITY

Cell line raw data generated in this study have been deposited to GEO. Patient raw data generated in this study have been deposited to EGA. The accession codes will be made available upon publication.

## MATERIALS AND METHODS

### Cell lines

Human CA46 (ATCC, #CRL-1648), HEL (ATCC, #TIB-180), SET-2 (DSMZ, #ACC 608) and CCRF-CEM (ATCC, #CRM-CCL-119) cell lines were maintained according to standard procedures in RPMI-1640 (Thermo Fisher Scientific, #11-875-119) with 10% (or 20% for CA46 and SET-2 cells) FBS (Thermo Fisher Scientific, #10-437-028) at 37°C with 5% CO_2_. Cell lines in culture were screened biweekly for mycoplasma contamination using the MycoAlert PLUS Mycoplasma Detection Kit (Lonza, #LT07-703).

### Patient samples

This study was approved by the local ethics committee and by the Institutional Review Boards of Weill Cornell Medicine, Memorial Sloan Kettering Cancer Center, and the Icahn School of Medicine at Mount Sinai, and was conducted in accordance with the Declaration of Helsinki protocol. Either fresh peripheral blood or cryopreserved mononuclear cells isolated from bone marrow biopsies or peripheral blood from patients with *JAK2^V617F^* mutations were retrieved after a database search (**Extended Data Table 1**). Samples were selected to harbor only *JAK2^V617F^* mutations, as assessed by MSK-IMPACT, RainDance, or Neogenomics hematology panels. Fresh peripheral blood mononuclear cells were isolated within 48 hours of blood collection utilizing a Ficoll (Thermo Fisher Scientific, #45-001-750) gradient according to manufacturer’s recommendations, and either processed immediately or cryopreserved for future experiments. Isolated mononuclear cells from peripheral blood or bone marrow biopsies were thawed if cryopreserved and stained according to standard procedures, beginning with resuspension in staining buffer (Biolegend, #420201) and incubation with Human TruStain FxC (10 minutes at 4°C; Biolegend, #422302) to block Fc receptor-mediated binding. Cells were then stained with a CD34-PE-Vio770 antibody (20 minutes at 4°C; Miltenyi Biotec, clone AC136, #130-113-180) and DAPI (Invitrogen, #D1306). The samples were then sorted for DAPI-negative, CD34-positive cells using a BD Influx cell sorter.

### Single nucleus ATAC-seq with GoT-ChA

Cells were subjected to nuclei isolation according to the Nuclei Isolation for Single Cell ATAC Sequencing protocol (version CG000169 Rev D, 10x Genomics). Briefly, cells were resuspended with lysis buffer (10 mM Tris-HCL (pH 7.4), 10 mM NaCl, 3 mM MgCl_2_, 0.1% Tween-20, 0.1% Nonidet P40 Substitute, 0.01% Digitonin, 1% BSA) and incubated on ice (3 minutes for patient samples, 5 minutes for cell_2_, 1% BSA, 0.1% Tween-20) and centrifuging to pellet isolated nuclei. Nuclei were then resuspended in 1X diluted nuclei buffer (10x Genomics) and counted using trypan blue and a Countess II FL Automated Cell Counter.

Nuclei were subsequently processed according to the Chromium Next GEM Single Cell ATAC Solution user guide (version CG000209 Rev F, 10x Genomics) with the following modifications:

1. During the GEM Generation and Barcoding reaction (Step 2.1), 1 μL of 22.5 μM GoT-ChA primer mix was added to the barcoding reaction mixture. The primers used are GoT-ChA_R1N (locus-specific primer with a Read 1N handle sequence: TCGTCGGCAGCGTCAGATGTGTATAAGAGACAG - [22bp locus specific]) and GoT-ChA_IS2 (locus-specific primer with an IS2 handle sequence: AGCAAGTGAGAAGCATCGTGTC - [22bp locus specific]). These primers allow for exponential amplification of the GoT-ChA fragments, relative to the linear amplification of ATAC fragments.
2. During the Post GEM Incubation Cleanup (Step 3.2), 45.5 μL of Elution Solution I is used to elute material from SPRIselect beads. 5 μL is used for GoT-ChA library construction, and the remaining 40 μL are used for ATAC library construction as indicated in the standard protocol.
3. To generate the GoT-ChA library, two additional PCRs were performed on the 5 μL set aside during Step 3.2 (**Extended Data Fig. 1c**). The first PCR aims to amplify genotyping fragments prior to sample indexing and uses P5 (binds the P5 Illumina sequencing handle: AATGATACGGCGACCACCGAGATCTACAC) and GoT-ChA_nested (a nested, biotinylated, locus specific primer with a TruSeq Small RNA Read 2 handle: /5BiosG/CCTTGGCACCCGAGAATTCCA-[22bp locus specific]) primers with the following thermocycler program: 95 °C for 3 min; 15 cycles of 95 °C for 20 s, 65°C for 30 s and 72°C for 20 s; followed by 72°C for 5 min and ending with hold at 4°C. After a 1.2X SPRIselect clean up, biotinylated PCR product is bound and isolated using Dynabeads M-280 Streptavidin magnetic beads (Thermo Fisher Scientific, #11206D). Briefly, beads are washed three times with 1X sodium chloride-sodium phosphate-EDTA buffer (SSPE, VWR, #VWRV0810-4L), added to the purified PCR product, and incubated at room temperature for 15 minutes. The beads are then washed twice with 1X SSPE buffer and once with 10 mM Tris-HCl (pH 8.0) before resuspending in water. The bead-bound fragments are then amplified and sample indexed using P5 and RPI-X (binds the TruSeq Small RNA Read 2 handle and adds a sample index and P7 Illumina sequencing handle: CAAGCAGAAGACGGCATACGAGATXXXXXXXXGTGAC TGGAGTTCCTTGGCACCCGAGAATTCCA, “X” denotes user-defined sample index) primers with the following thermocycler program: 95 °C for 3 min; 6-10 cycles of 95 °C for 20 s, 65°C for 30 s and 72°C for 20 s; followed by 72°C for 5 min and ending with hold at 4°C.

Final libraries were quantified using a Qubit dsDNA HS Assay Kit (Thermo Fisher Scientific, #Q32854) and a High Sensitivity DNA chip (Agilent Technologies, #5067-4626) run on a Bioanalyzer 2100 system (Agilent Technologies) and sequenced on a NovaSeq 6000 at the Weill Cornell Medicine Genomics Resources Core Facility with the following parameters: paired-end 50 cycles; Read 1N 50 cycles, i7 Index 8 cycles, i5 Index 16 cycles, Read 2N 50 cycles. ATAC libraries were sequenced to a depth of 25,000 read pairs per nucleus and GoT-ChA libraries were sequenced to 5,000 read pairs per nucleus.

### Single cell ASAP-seq with GoT-ChA

Samples were processed in a similar fashion to that described previously for standard scATAC-seq, with a few key differences as described by the original authors^38^. Additional minor modifications for incorporation of GoT-ChA into ASAP-seq are as follows:

1. During the GEM Generation and Barcoding reaction (Step 2.1), 1 μL of 22.5 μM GoT-ChA primer mix is added to the barcoding reaction mixture, just as was done for scATAC-seq.
2. During the Post GEM Incubation Cleanup (Step 3.2), 45.5 μL of Elution Solution I is used to elute material from SPRIselect beads. 5 μL is used for GoT-ChA library construction, and the remaining 40 μL are used for ATAC library construction as indicated in the standard protocol. GoT-ChA library construction proceeded as described above.

Final libraries were quantified using a Qubit dsDNA HS Assay Kit (Thermo Fisher Scientific, #Q32854) and a High Sensitivity DNA chip (Agilent Technologies, #5067-4626) run on a Bioanalyzer 2100 system (Agilent Technologies) and sequenced on a NovaSeq 6000 with the following parameters: paired-end 50 cycles; Read 1N 50 cycles, i7 Index 8 cycles, i5 Index 16 cycles, Read 2N 50 cycles. ATAC libraries were sequenced to a depth of 25,000 read pairs per cell, and both GoT-ChA and any protein tag libraries were sequenced to 5,000 read pairs per cell.

Protein expression measurements were performed using TotalSeq-A reagents from Biolegend according to manufacturer’s recommendations. The following surface markers were assayed: CD34 (#343537), CD38 (#356635), CD90 (#328135), CD49f (#313633), CD45RA (#304157), CD41 (#303737), CD36 (#336225), CD69 (#310947), CD9 (#312119), CD71 (#334123), CD99 (#371317), CD184 (#306531), HLA-DR (#307659), CD134 (#350033), CD48 (#336709), CD52 (#316017), CD135 (#313317), CD47 (#323129), CD7 (#343123), CD56 (#362557), and CD45 (#368543).

### GoT-ChA data processing

#### Initial read processing and genotype calling

Raw sequencing data for GoT-ChA libraries were demultiplexed using cellranger-ATAC mkfastq. The GoT-ChA sequencing FASTQ files were then used as input into the GoT-ChA pre-processing pipeline designed to result in a genotype-per-cell output. These functions are available as a R package (“Gotcha”; see code availability section).

As a first step, input FASTQ files are split into a user-defined *n* reads files to allow for parallelized downstream processing through the “FastqSplit” function. Next, the “FastqFiltering” function takes each newly generated split FASTQ file and identifies read pairs that do not pass a set of user-defined parameters for base quality filtering. Default usage was designed to identify poor quality bases at or surrounding a SNV site, though the function includes parameters to easily adjust for filtering of all base pairs in paired read sequences or of only a single read for each pair. After quality filtering, the “BatchMutationCalling” function first identifies whether a read contains the expected sequence of the nested primer used during library construction through pattern matching. If so, then each read’s paired cell barcode is matched to a provided whitelist, considering a maximum Hamming distance of two. All reads that pass these criteria are then assessed for whether the read contains a specified wildtype or mutant sequence at a designated position. This process is performed for each read in each split and filtered FASTQ file via parallelized processing through a slurm workload manager, ultimately outputting a cell barcode by genotyped reads matrix for each FASTQ file processed. Finally, “MergeMutationCalling” combines all matrices generated from each split FASTQ file and merges them together grouping by cell barcode and summarizing the counts of reads that were identified as wildtype, mutant, or neither. The summarized genotyping data must then be integrated with the chromatin accessibility information via shared cell barcodes. This is achieved via the “AddGenotypingArchR” function, that is compatible with the ArchR^157^ scATAC-seq pipeline.

#### Background correction and genotype assignment

Once read counts per cell corresponding to the WT and MUT alleles are generated, we then require a method to label these cells with their corresponding biological genotype. Given high level of PCR amplification needed to capture in single cells the 1-2 molecules of DNA containing the targeted locus, the data is vulnerable to various sources of noise such as PCR errors and PCR recombination, aggravated by the exponential amplification bias inherent to PCR. For example, the total observed read counts for WT do not correspond to the number of WT alleles. Rather, they are defined by the cycle of PCR in which the targeted loci is captured, as well as the sequencing depth. Thus, if an allele is captured earlier on in the PCR process, exponential amplification would then result in inflated read counts. Additionally, ambient contamination of fragments between neighboring cells can influence the null distribution of reads for each allele. To account for the potential sources of noise, we developed two alternative approaches to quantify and correct for potential background noise in the genotyping data. Our first approach (*empty droplet-based*) leverages the genotyping information present in empty droplets generated during the 10X run (i.e., barcodes that do not contain a cell, and therefore yield low to no ATAC-seq fragments). Our second approach, (*cluster-based*) leverages the presence of a bimodal distribution of genotyping reads in cells, representing successful vs. unsuccessful genotyping. Both methods are described in the sections below.

#### Empty droplet-based background noise correction and genotype assignment

To estimate the background noise present in the genotyping data, we leveraged the presence of empty droplets obtained from in every 10X run, as has previously been done for noise correction in single cell protein expression^158^. First, background noise is estimated for either wildtype or mutant reads independently. Given the zero-inflated distribution of genotyping reads present in empty droplets and to avoid the potential presence of outlier values (i.e. a droplet that contains a cell but was assigned as empty), we estimate the value of the background noise as that of the 99^th^ percentile of the read number distribution for each genotype independently. Once the background noise is quantified, we proceed to subtract the value for each genotype read count from the barcodes containing real cells. In addition, cells are required to contain a minimum number of genotyping reads (>250 after background subtraction). This procedure can be performed by using the “AddGenotyping” function followed by the “FilterGenotyping” function, both available in the Gotcha package (**see code availability**).

#### Cluster-based background noise correction and genotype assignment

An alternative approach that results in higher genotyping efficiency and includes the ability to detect heterozygous mutated cells, was informed by pre-existing approaches for normalization of CITE-seq data and hashtag oligo (HTO) demultiplexing such as DSB^158^ and HTODemux^159^, respectively. The basic assumption is that in our dataset we may encounter populations of homozygous WT, homozygous MUT, and heterozygous cells. In addition, we anticipate a population of cells for which no genotyping call can be made, due to experimental constraints on capture. These cells may still have non-zero read counts for each allele, representing the level of background signal for each allele. Thus, the data will typically follow a bimodal distribution of supporting reads for either the mutated or wildtype allele. One mode reflects cells with true capture of the mutated allele and the second mode reflects cells where reads reflect background noise.

In the DSB^158^ method, the authors employ a log-transformation of the protein expression counts, which produces a bimodal distribution where the lower mode represents the background signal and the higher mode represents the true signal. The log-counts are then z-scored using the mean and standard deviation of the noise distribution, as defined by a Gaussian Mixture Model (GMM) with 2 components. In several methods for HTO demultiplexing, the transformed hashtag counts are clustered, and the resulting clusters are characterized to produce interpretable labels. Therefore, we sought to combine these two basic steps: normalization based on background noise distribution followed by clustering.

The flowchart depicting the steps of our method is shown in **Extended Data Fig. 2f**. All analyses, including estimation of background, is performed only on the set of barcodes representing captured cells as defined by the ATAC library profile (see section **ATAC-seq data processing** below). Each feature (WT and MUT read counts) is normalized independently. First, the read counts are log-transformed with a pseudocount of 1. This results in a bimodal distribution as shown in **Extended Data Fig. 2g**. However, since count data is discrete, continuous density estimation methods such as GMMs and Kernel Density Estimation (KDE) are not immediately suitable. Thus, we add a Gaussian-distributed, independent smoothing noise with mean 0 and standard deviation 0.3. This results in a continuous, bimodal distribution of transformed counts, as shown in **Extended Data Fig. 2g**. This smoothing noise is later removed (see below).

Next, we define a probability density function (PDF) on this data for decomposition background signal and true signal distributions. For this purpose, we use a KDE with a Gaussian kernel and a fixed bandwidth. The optimal bandwidth is inferred through a cross-validation grid search; for each possible bandwidth value between 0.1 and 1.0, we randomly select 10% of the data and compute the KDE using these values as input. We then find the log-likelihood of this model on the entire dataset. This process is repeated 10 times for each bandwidth value, and the “loss” is defined as −1*mean(cross-validated log-likelihood). This loss is plotted vs. each bandwidth value, and a polynomial is fit to approximate the loss function. This function is then minimized using stochastic gradient descent to find the optimal bandwidth parameter.

Next, the value defining the boundary between background and signal distributions is defined as the lowest point in the KDE between the two modes. The KDE is decomposed by computing KDEs separately on the data under or above the defined boundary, where the weights of each PDF component are defined by the proportion of the data below or above the threshold. Then, the mean and standard deviation of the inferred distribution of the measurement background are computed by integration. The effect of the Gaussian smoothing noise is accounted for as follows. The KDE we have just modeled is the PDF of the random variable X+Y. Then, given the properties of expectations (E) and variances (VAR), where X is the raw log-transformed read counts and Y = N(0,0.09) is the smoothing noise:

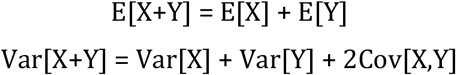

Since X and Y are independent, the covariance term is zero. To compute z-scores, we then go back to the raw, log-transformed counts, and subtract the mean and divide by the standard deviation of the background noise distribution estimated from the PDF of X+Y, adjusted for Y. This is a simple subtraction of Var[Y] = 0.09 from the calculated variance. Importantly, as opposed to the GMM approach, this approach does not force the inferred background noise and signal distributions to be Gaussian. In fact, the observed background noise across multiple samples shows a positive skewed distribution. Thus, this approach is more broadly generalizable to unseen datasets. Below is the formulation of the final z-score:

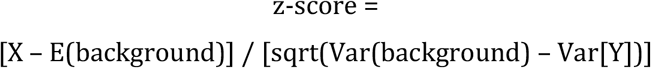

Once again, X represents the log transformation of the read counts, with pseudocount 1. The distribution of the background noise is computed as the KDE of the values of X+Y below the previously computed threshold. We can then plot the z-scores for each WT and MUT distributions in two dimensions (wildtype z-scores and mutated z-scores), with the thresholds dividing the data into four quadrants as shown in **Extended Data Fig. 2h**. We set the minimum z-score threshold in each dimension to be 2.0 to account for potential edge cases in the upstream calculation. The resulting quadrants are then easily interpretable: cells in the lower left quadrant have low z-scores for both WT and MUT and as such are non-genotyped. The lower right quadrant corresponds to homozygous WT cells, the top left is homozygous MUT, and the top right consists of heterozygous cells. Crude genotype calls can in fact be made at this stage and these labels are included in the final output. However, since decision boundaries are rarely optimal, we proceeded with a clustering-based approach, based on spectral clustering and KDE-based mixture models. The underlying intuition for this step is that the clustering method can be further informed by the joint distribution of the data for WT and MUT z-scores. Thus, we again fit a KDE to the data, this time in two dimensions, while again adding a small multivariate Gaussian noise for smoothing and better fit. The optimal bandwidth was computed using the procedure described above. This bandwidth was then used as the hyperparameter for spectral clustering of the data using a Gaussian kernel. To choose the parameter “k” for the number of clusters, we restricted our search to between 2 and 4 clusters for interpretability, where one cluster will always represent non-genotyped cells. The optimal “k” was chosen by computing the Adjusted Rand Index between each clustering output and the quadrant labels and picking the best corresponding “k”, with the rationale being that clusters should line up with the quadrant divisions for biological interpretability. Clusters were characterized by the quadrant in which their marginal median lie (i.e., the vector corresponding to the medians along each feature).

Once the clusters were defined on the smoothed data, we then used them to generate a KDE mixture model. KDE was computed using the previously computed optimal bandwidth on each cluster separately, with mixture weights being initialized as the cluster proportions and updated through expectation maximization. This generated a probabilistic model that we could then use to compute the probability of every cell belonging to each genotype cluster. These probabilities can be computed on the raw z-scores without smoothing Gaussian noise added during the initial KDE fits. Thus, we now have relative likelihoods of each cell belonging to each identified genotype, and hard cluster assignments could be created by assigning cells to their most likely cluster.

However, since this model was produced from the smoothed data, we next proceed to refine the labeling. To do this, we applied techniques from semi-supervised learning. Since the KDE mixture model (learned from the Z-scores with 2D Gaussian noise added) was likely to produce confident labels when the maximum assigned probability is high, we elected to remove the labels of all the cells that had a probability of belonging to their assigned genotype below 99%. These unlabeled points tended to lie at the cluster boundaries. To relabel them, we used a semi-supervised K-Nearest Neighbors classifier. With k = sqrt(N), the classifier would build a KNN graph using all the data and iteratively assign unlabeled points based on the labels of their neighbors. The edges of the KNN graph were weighted using an RBF kernel to account for the distance between neighbors. Given the size of most single-cell datasets, we chose to opt for precision over performance, and forced the classifier to re-label only the most optimal point each time before the KNN graph is re-computed on each iteration.

In this way, all the previously unlabeled points are labeled by their closest inferred genotype. The pipeline outputs and stores all relevant plots for each sample. It is recommended for the user to inspect plots to assess the two possible outputs (quadrant vs. cluster-based genotyping) and determine which output’s labels fit the distribution of the data better. The default choice should be the cluster outputs, except in the case the assigned clusters span multiple quadrants to a large degree. Finally, the output data is stored for cell barcode matched integration into the scATAC-seq metadata using the “AddGenotype” function for downstream analysis. The genotype labeling analysis was performed in Python, using the packages pandas, numpy, matplotlib, seaborn, scipy, and sklearn. Detailed documentation of the method and plots at each step are available (**see code availability**).

### ATAC-seq data processing

#### Cell line mixing data processing and analysis

Raw sequencing data for ATAC-seq libraries were demultiplexed using cellranger-ATAC (v2.0.0) mkfastq. ATAC sequencing reads were then aligned to the hg38 reference genome using cellranger-ATAC count. Fragment files generated by cellranger-ATAC were used as input for processing through the ArchR (v1.0.1) pipeline^157^ for downstream analysis. Based on barcode quality control, a minimum TSS enrichment score of 5 and a minimum number of unique fragments of 20,000 was set based on data distribution. Potential doublets were identified by the addDoubletScores function and removed using the filterDoublets function. Initial dimensionality reduction was performed via iterative latent semantic indexing (LSI) using the cell by genomic bin matrix (bin size = 500 bp) with the following parameters: iterations = 4, resolution = 0.1-4, sampleCells = 10,000, n.start = 10. Cell clustering was performed through the addClusters function using the Seurat method, with a resolution of 0.5.

#### Patient data processing dimensionality reduction and clustering

Raw sequencing data for ATAC-seq libraries were demultiplexed using cellranger-ATAC (v2.0.0) mkfastq. ATAC sequencing reads were then aligned to the hg38 reference genome using cellranger-ATAC count function. Fragment files generated by cellranger-ATAC were used as input for the ArchR (v1.0.0) pipeline^157^ to generate the cell by genomic bin matrices (bin size = 500 bp) for each patient sample. High quality barcodes representing captured cells were identified based on TSS enrichment score above 7.5 and a minimum number of unique fragments above 3,000 based on their distribution as shown in **Extended Data Fig. 3b**. For initial dimensionality reduction, the cell by genomic bin matrix was used as input for reciprocal latent semantic indexing (LSI) as calculated by the Signac (v1.1.1) pipeline^52^. Briefly, the term frequency-inverse document frequency (TF-IDF) matrix is calculated followed by singular value decomposition as implemented in the RunSVD function. Dimensionality reduction via UMAP was performed using the RunUMAP function of the Seurat (v4.0.1) package^160^, for LSI components 1:50. Next, to generate an integrated embedding across patient samples, we performed unsupervised identification of anchor correspondence between datasets^161^ using the “FindIntegrationAnchors” function. To generate a common LSI space, we ran the IntegrateEmbeddings^161^ function followed by the RunUMAP function with the following parameters: dims = 1:50, min.dist = 0.01 and spread = 0.5 for dimensionality reduction for visualization in two dimensions. Cell clustering was performed using the “FindNeighbours” function using the first 50 dimensions of the integrated LSI space as input followed by the “FindClusters” function with resolution = 1. Downstream analysis was performed based on the genotype assignment obtained through the Gotcha pipeline (see GoT-ChA data processing section above). These calculations can be performed by running the “NormalizedMutantFraction” function of the Gotcha R package. Pseudotime cell ordering was performed in a semi-supervised manner as described in the ArchR (v1.0.0) pipeline^157^. For calculating the fraction of mutant cells along pseudotime as defined by the differentiation axis (erythroid, megakaryocytic, neutrophil-monocytic or lymphoid), cells involved in the differentiation trajectory were divided into 10 quantiles based on their pseudotime values, and the fraction of mutant cells within the window was calculated for each sample.

#### Differential gene and motif accessibility modeling inter-patient variability

Gene accessibility scores were obtained through the ArchR (v1.0.0) pipeline^157^, and ChromVAR^70^ (v1.8.0) was utilized within the ArchR pipeline to estimate transcription factor motif accessibility z-scores. For intra-cluster differential testing of either gene or transcription factor motif accessibility, we took a statistical approach allowing to account for potential technical confounders arising from sample-specific batch effects. To that end, we applied a linear mixture model (LMM) approach followed by likelihood ratio test as follows:

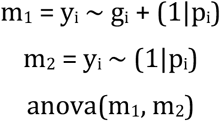

Where y_i_ is the response feature for cell i, represented by either motif or gene accessibility scores, g_i_ is the genotype of cell I and pi is the patient sample of origin for cell i explicitly model as a random factor of the LMM model. This model was selected to account for patient specific confounders, and to account for variability in cell numbers across samples. Raw P values were then adjusted by Benjamini-Hochberg correction. This analysis can be applied using parallelized computing via the “DiffLMM” function available within the Gotcha R package (see **code availability** section). For gene accessibility scores, accessibility differences were calculated as the ratio of the mean value across the cells for the specified genotype and cell cluster for mutant versus wildtype cells, followed by application of log_2_. For transcription factor motif accessibility, differences between genotypes were calculated as the difference in the mean z-score for each transcription factor for the specified cell cluster and genotype.

Pathway enrichment analysis was performed via preranked differential gene accessibility scores using the msigdbr (v7.2.1) and the fgsea (v1.12.0) packages. The differential gene accessibility rank was generated by utilizing the −log_10_(FDR) times the sign based on the direction of change (1 or −1). This rank was then used as input for a pre-selected set of Hallmark pathways into the fgsea function with minSize = 10, maxSize = 1,000 and nperm = 100,000.

Transcription factor motif accessibility correlations were calculated based on the z-scores matrices estimated by ChromVAR^70^ (v1.8.0). Cell by motif matrices were subset based on the transcription factors of interest and motif-motif correlations were calculated via the cor function in R (v3.6.2). For STAT1 and NFKB1 correlations, the linear model was estimated using the “lm” base function in R, and F-test parameters were retrieved using the “summary” base R function.

Co-accessibility scores for either wildtype or mutant cells for a given cluster were calculated leveraging the ArchR “addCoAccess” function based on Cicero^123^. We generated a wrapper function named “DiffCoAccessibility”, allowing to split co-accessibility calculations by genotype, generating loop files for each group separately. Briefly, after specifying the cell clusters, genotype levels (i.e., WT, MUT or HET), subset of peaks of interest and the maximum distance between pairs of peaks, co-accessibility is calculated for each genotype independently, and the loop files for each genotype are provided as output. The output of this function can then be provided to the “plotDCA” function to visualize the co-accessibility loops simultaneously for both genotypes for each peak. Both functions are available as part of the Gotcha R package.

### Protein data processing

Protein expression was estimated using the antibody derived tag information. ADT FASTQ files were first processed with a python script “ASAP_to_kite.py” (obtained from ASAP-seq authors) which converts the files into a format similar to the 10x scRNA-seq FASTQ format, thus enabling analysis using the kallisto^162^ (v0.46.0), bustools^163^ (v0.39.3), and kite^164^ (v0.0.2) frameworks. Protein data was normalized via the “DSBNormalizeProtein” function from the DBS (v0.1.0) package^158^. Statistical comparisons between genotypes were performed via Wilcoxon rank sum test followed by Bonferroni multiple hypothesis correction.

### Mitochondrial data pre-processing and variant calling

A modified version of the hg38 reference genome with hard-masked nuclear mitochondrial DNA sequences (NuMTs) was generated using cellranger-atac (v2.0.0) “mkref” function. Sequencing reads were then mapped to the modified reference using cellranger-atac count. Note that most scATAC fragments are not expected to arise from NuMTs, so they can be a priori safely uniquely mapped to the mitochondrial genome by masking their nuclear paralogs^37^. Next, the mgatk (v0.5.0) package function “mgatk” with “tenx” mode was used to generate base counts at each mitochondrial genomic position for each cell passing cell-ranger quality control standards. Reads with a mapping quality lower than 20 were not considered, as well as bases with a sequencing quality lower than 20. Files containing the combined per-cell base count information for each sample were loaded into R and analyzed using custom code and functions developed to identify mitochondrial variants^165^. In summary, only those variants supported by at least 2 forward and 2 reverse reads in a minimum of 5 cells, likely heteroplasmic (variance to mean ratio in log scale higher than −2) and with a high strand concordance (correlation between forward and reverse counts across cells higher than 0.65) were considered for downstream analysis^37^. Some previously described false-positive calls were further discarded, as well as those variants present in two or more unrelated samples, since it points out to arise from technical artifacts. For each variant call, an alternative allele frequency—also called heteroplasmy in mitochondrial context—was calculated for each cell. However, for those cells showing less than 10 reads the allele frequency was set to undetermined to favor robust calling of heteroplasmic fractions. The GoT-ChA genotyping information was combined with these results and used to visualize mitochondrial mutations present in a subgroup or the whole population of cells carrying the GoT-ChA-targeted mutation. In this situation, both the nuclear and mitochondrial mutations are referred to as being in phase.

### Cell genotype classifier based on mitochondrial variants

Mitochondrial mutations in phase with the GoT-ChA mutation status were used to impute genotyping of cells whose mutation status could not be determined based on GoT-ChA information. This was achieved by implementing a supervised learning approach based on a random forest classifier. For training and testing the classification model, the dataset was downsampled to contain the same number of mutant and wildtype cells to obtain a balanced dataset for training and test, and 90% of the cells were assigned to the training set and 10% were assigned to the test set. The classification model was built by applying the “randomForest” function within the R package tidymodels (v0.1.3)^166^ to the training set, using the heteroplasmy of the in-phase mutations as features and wildtype and mutant genotypes as classes. The number of decision trees was set to 1000. The model was then applied to the test set to obtain genotype predictions. The classifier accuracy was measured as the percentage of correctly classified instances out of all predictions. As the model showed a high accuracy on the test set, it was subsequently applied to make genotype imputations for the undetermined GoT-ChA calls.

### Inference of copy number variations (CNV) from scATAC-seq data

A CNV score was calculated for each cell adapted from a method previously described^36^. Here, instead of using bins of similar GC content as a baseline CNV score, we compared the normalized counts in 10 Mb bins (step size of 2 Mb) to normal diploid CD34^+^ cells. CNV scores were plotted as a heatmap for visualization and hierarchical clustering was performed using fastcluster (v1.2.3) and parallelDist (v0.2.6) R packages using default parameters.

### Cell label prediction based on scRNA-seq reference and bridge integration

To orthogonally validate our manually assigned cluster labels, we leveraged a novel method for annotation of scATAC-seq cell clusters using a scRNA-seq reference dataset (available at https://zenodo.org/record/5521512#.YmnDDi-B1uV), via multiome (scATAC-seq plus scRNA-seq) bridge integration through Seurat (v4.1.0)^160^. Briefly, genomic features present in the publicly available multiome data (GSE194122) were utilized to recalculate the count matrices from the scATAC-seq query data. A common dimensionality reduction space was generated allowing for the creation of a bridge reference, followed by defining bridge anchors between the extended bridge reference and the scATAC-seq query. Finally, cells from the query were mapped to the reference cluster labels via the extended bridge integration.

### *JAK2^V617F^* reversible mouse model, RNA-seq, and data analysis

*Jak2^Rox/Lox^* (*Jak2^RL^*) *Dre-rox, Cre-lox* dual recombinase knock-in/knock-out mice^110^ were crossed to UbcCreER tamoxifen-inducible Cre lines and RLTG dual-recombinase reporter lines^167,168^. *Jak2^V617F^* knock-in was carried out using Dre mRNA electroporation^110^. For gene expression analysis, secondary cohorts of lethally irradiated C57BL/6mice transplanted with UbcCreER-*Jak2^RL^* bone marrow 8 weeks post-transplant and exhibiting MPN were treated with ruxolitinib (60 mg/kg P.O. BID), tamoxifen (100 mg/kg daily x4 followed by 80 mg/kg daily of TAM chow x3) or vehicle (MPN control) for 7 days and then sacrificed. Bone marrow cells were isolated from limb bones into FACS buffer (phosphate buffered saline (PBS) + 2% fetal bovine serum) via centrifugation (8000 rpm x 1 min). After red blood cell lysis (BioLegend), single-cell suspensions were depleted of lineage-committed hematopoietic cells using a Lineage Cell Depletion Kit according to manufacturer’s protocol (EasySep™, StemCell Technologies, Inc.). Lineage-depleted bone marrow was then stained with an antibody cocktail comprised of a lineage cocktail (BioLegend), as well as c-Kit (2B8), Sca-1 (D7), FcγRII/III (2.4G2) and CD34 (RAM34). All antibodies were purchased from BioLegend. After staining, samples were washed in FACS buffer and resuspended in FACS buffer with DAPI as a live-dead stain for cell sorting. TdTomato+ (*Jak2^RL^* knock-in) or GFP+ (*Jak2^RL^* knockout) LSKs and MEPs were then sorted on a FACSAria III directly into Trizol LS (Invitrogen) and stored at −80°C until processing. RNA was subsequently isolated using the Direct-Zol Microprep Kit (Zymo Research, R2061) according to manufacturer’s protocol and quantified using the Agilent High Sensitivity RNA ScreenTape (Agilent 5067-5579) on an Agilent 2200 TapeStation. cDNA was generated from 1ng of input RNA using the SMART-Seq HT Kit (Takara 634455) at half reaction volume followed by Nextera XT (Illumina FC-131-1024) library preparation. cDNA and tagmented libraries were quantified using High Sensitivity D5000 ScreenTape (5067-5592) and High Sensitivity D1000 ScreenTape respectively (5067-5584). Libraries were sequenced on a NovaSeq at the Integrated Genomics Operation (IGO) at MSKCC.

## EXTENDED DATA

**Extended Data Fig. 1.**
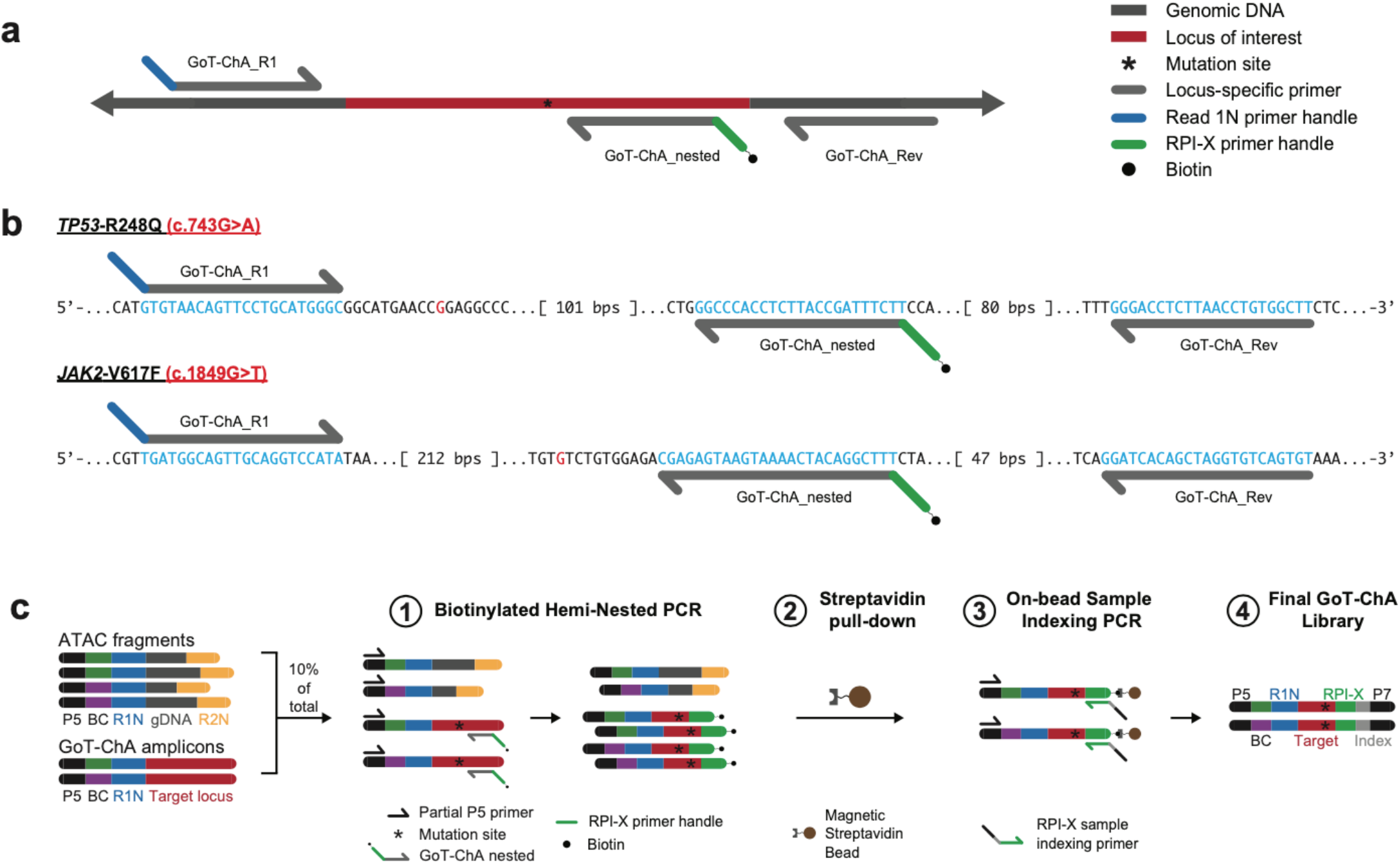
Primer sequences, relative positions, and library construction schematic for GoT-ChA. **a**, Primer design schematic for GoT-ChA, indicating the relative position of GoT-ChA primers to one another as well as the targeted locus of interest. **b**, Primer binding sites for *TP53^R248^* and *JAK2^V617^* genotyping. Binding sites are highlighted in blue, with custom primer handles shown as indicated in panel **a**. **c**, Schematic representation detailing GoT-ChA library construction, comprised of a biotinylated hemi-nested PCR, a streptavidin-biotin pull-down, and an on-bead sample indexing PCR, ultimately resulting in the generation of a genotyping library compatible with Illumina sequencing. P5. Illumina sequencing handle; BC, unique cell barcode; R1N, Read 1 Nextera adapter; gDNA, genomic DNA; R2N, Read 2 Nextera adapter.

**Extended Data Fig. 2.**
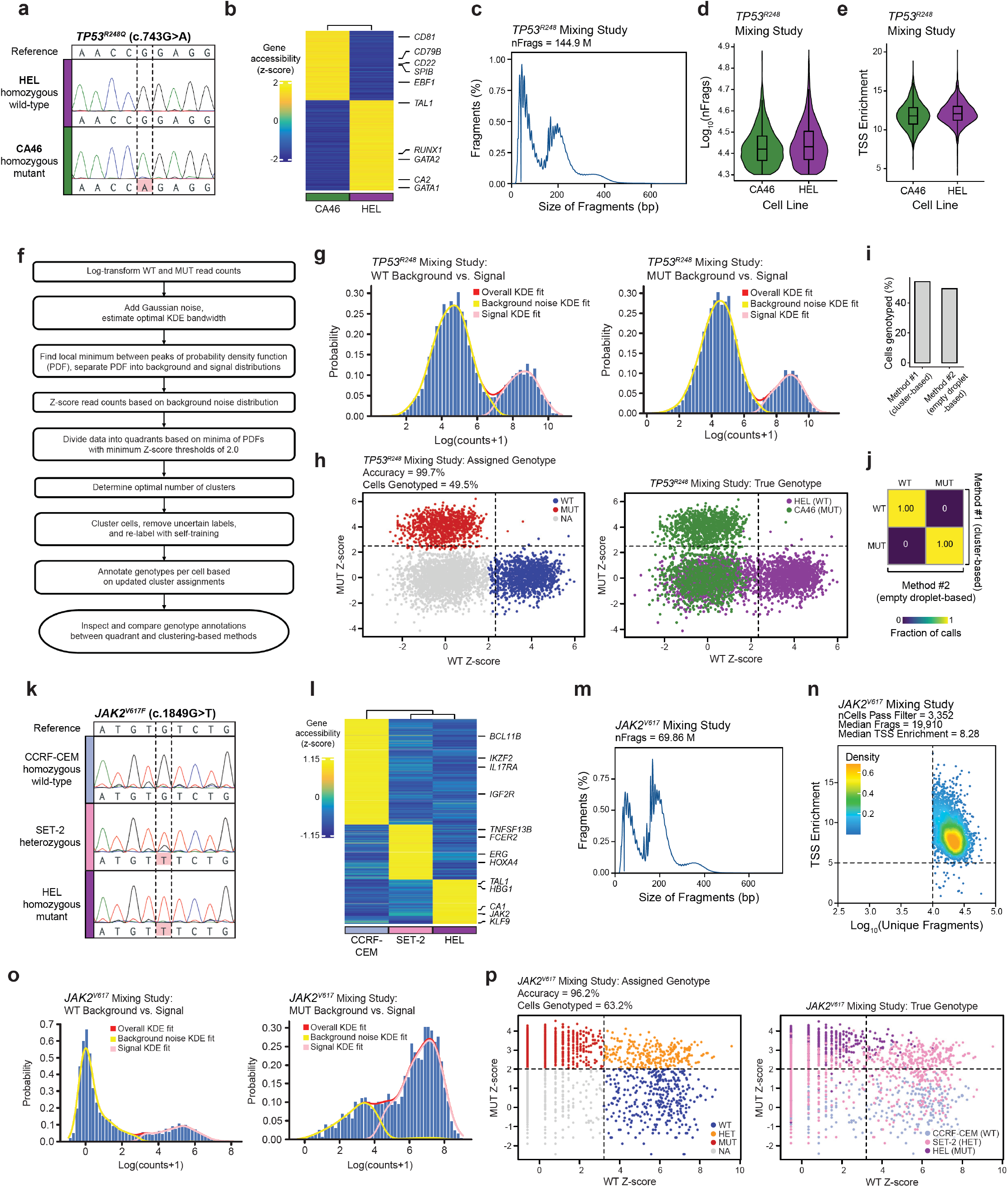
Genotyping accuracy, quality control metrics, and GoT-ChA data processing of proof-of-principle mixing studies. **a**, Sanger sequencing confirming known homozygosity of *TP53^R248^* wildtype HEL cells and *TP53^R248Q^* mutant CA46 cells. **b**, Differential gene accessibility score heat map illustrating the identification of the two human cell lines HEL and CA46 in the *TP53^R248^* mixing study (FDR < 0.05 and log_2_FC > 1.25; Wilcoxon rank sum test followed by Benjamini-Hochberg correction). **c**, scATAC-seq library fragment size distribution for the *TP53^R248^* mixing study, illustrating characteristic periodicity due to nucleosome positioning. **d**, Number of unique nuclear fragments per cell for each cell line in the *TP53^R248^* mixing study, indicating adequate complexity of the scATAC-seq libraries. **e**, TSS enrichment scores per cell for each cell line in the *TP53^R248^* mixing study, indicating a high signal-to-background ratio in the scATAC-seq data. **f**, Schematic representation illustrating the workflow of the analytical pipeline used for noise correction of the GoT-ChA data and for defining single cell genotype assignments. **g**, Histograms of WT (left panel) and MUT (right panel) number of reads per cell from the *TP53^R248^* mixing study. Kernel density estimation (KDE) lines for overall data (red), background (yellow), and signal (pink) are shown for each genotype. **h**, Scatter plots comparing assigned genotypes from GoT-ChA processing (left panel) as compared to the true genotypes as determined by cell line identity (right panel). Dotted lines indicate the detected threshold for the distinction between background and signal before updated cluster assignments for both WT and MUT data. **i**, Comparison of the percentage of cells genotyped in alternative empty droplet-based noise correction methods (see materials and methods). **j**, Confusion matrix for genotype assignments for those cells with available genotyping in both alternative noise correction methods (see materials and methods). **k**, Sanger sequencing confirmation of known genotypes for the *JAK2^V617^* mixing study: CCRF-CEM wildtype cells, SET-2 heterozygous cells, and HEL homozygous mutant cells. Note that SET-2 data confirm the known allelic ratio of 3:1 for mutated:wildtype alleles in this cell line. **l**, Differential gene accessibility score (FDR < 0.05 and log_2_FC > 1.25; Wilcoxon rank sum test followed by Benjamini-Hochberg correction) heat map indicating identification of the three human cell lines CCRF-CEM, SET-2, and HEL used in the *JAK2^V617^* mixing study. **m**, Fragment size distribution for the *JAK2^V617^* mixing study scATAC-seq library, showing expected nucleosomal periodicity. **n**, Scatter plots showing the number of unique nuclear fragments per cell vs. the transcriptional start site (TSS) enrichment. Dotted lines indicate the selected thresholds based on the distribution. **o**, Histograms of WT (left) and MUT (right) read distributions from the *JAK2^V617^* mixing study. KDE lines for overall data (red), background (yellow), and signal (pink) are shown for each genotype. **p**, Scatter plots comparing assigned genotypes from GoTChA processing (left panel) compared to the true genotypes (right panel) as determined by cell line identity. Dotted lines indicate the initial thresholds identified between background noise and signal for either WT (vertical line) or MUT (horizontal line) data before final genotype assignment after clustering (see materials and methods).

**Extended Data Fig. 3.**
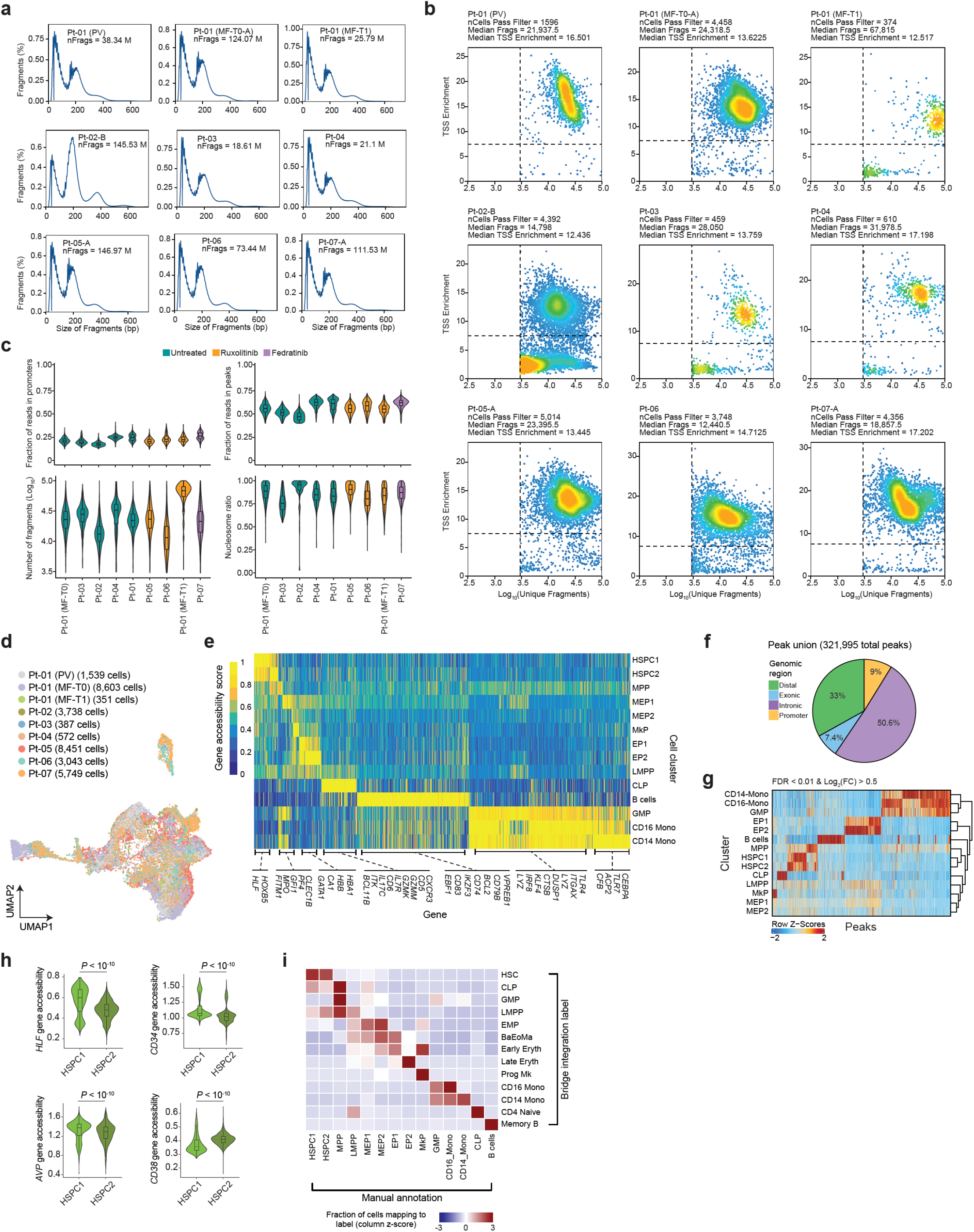
GoT-ChA results in high quality chromatin accessibility data in human patient samples, allowing progenitor subset annotation. **a**, Fragment size distribution plots per sample, showing characteristic periodicity due to nucleosome positioning across all samples. In samples for which multiple 10x lanes were run, a representative example is shown. **b**, Scatter plots showing the number of unique nuclear fragments versus TSS enrichment for each patient sample, with dotted lines indicating the thresholds used for identification of high-quality cells (TSS enrichment score > 8 and log_10_(unique fragments) > 3.5). The median values for unique nuclear fragments, median TSS enrichment score, and total number of cells that pass filters per patient sample are shown. **c**, Violin plots detailing quality metrics for scATAC-seq data, plotted per patient, and colored by JAK inhibitor therapy: fraction of reads in promoters (top left), fraction of reads in peaks (top right), number of unique fragments (bottom left), and nucleosome ratio (bottom right). Untreated samples are colored green, ruxolitinib-treated in yellow, and fedratinib-treated in purple. **d**, Integrated uniform manifold approximation and projection (UMAP) colored by patient sample, illustrating consistent distribution of cells after integration through reciprocal latent semantic indexing (see materials and methods). **e**, Heat map of gene accessibility scores across all progenitor subclusters, showing genes with differential accessibility (FDR < 0.05 and log_2_FC > 1; Wilcoxon rank sum test followed by Benjamini-Hochberg correction), and highlighting informative marker genes. **f**, Genomic distribution of peaks in distal regions (green), exonic regions (blue), intronic regions (purple), and promoters (yellow) for the union of peaks (n = 321,995 peaks). **g**, Differentially accessible peaks (FDR < 0.05 and log_2_FC > 1; Wilcoxon rank sum test followed by Benjamini-Hochberg correction) across progenitor clusters. **h**, Gene accessibility scores for key stem cell (*HLF, AVP, CD34, CD38*) marker genes across clusters HSPC1 and HSPC2 (Wilcoxon rank sum test), consistent with HSPC1 representing earlier HSPCs (higher *HLF, AVP* and *CD34*) and HSPC2 representing later HSPCs (higher *CD38* and lower *HLF, AVP* and *CD34*). **i**, Confusion matrix between manually annotated cluster labels and predicted labels based on scRNA-seq reference via bridge integration mapping^54^ (see materials and methods).

**Extended Data Fig. 4.**
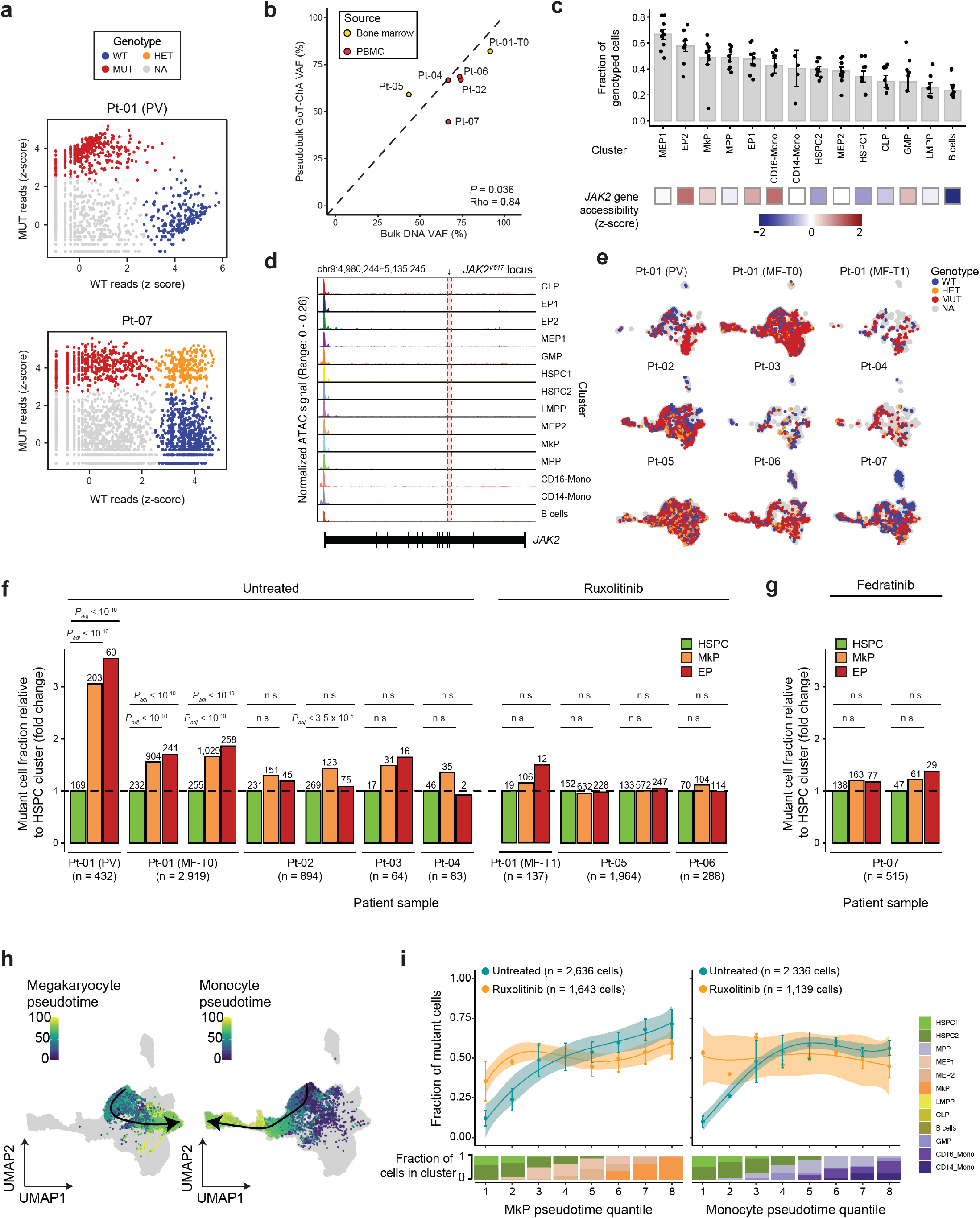
Genotyping and distribution of mutant cells in patient samples. **a**, Scatter plots illustrating two representative examples of assigned genotypes from GoT-ChA processing. Final genotype assignment after clustering shown in dot color. Homozygous wildtype (WT; blue), homozygous mutant (MUT; red), heterozygous (HET; yellow) or not assignable (NA; grey). **b**, Scatter plot of pseudobulk GoT-ChA variant allele fraction (VAF) versus bulk DNA sequencing VAF reported clinically for those patients for which was available (n = 6 patients; no clinically reported VAF available for samples Pt-01 (PV), Pt-03, and Pt-01 (MF-T1)). Source of biological material for each patient sample used for GoT-ChA (bone marrow in yellow and peripheral blood in red) is indicated. Dotted line represents a slope of 1 with intersection 0 as reference. Correlation values are shown (Spearman correlation test). **c**, Fraction of genotyped cells and average *JAK2* gene accessibility per cluster. Bars represent the mean; each point represents a patient sample and error bars represent the standard deviation (upper panel). *JAK2* mean gene accessibility z-scores across cell clusters is shown (bottom panel) **d**, Chromatin accessibility coverage of the *JAK2* gene across progenitor clusters. The V617 hotspot is indicated by the region shaded red. **e**, Genotype calls projected onto the integrated uniform manifold approximation and projection (UMAP) per sample. WT = wildtype cells (blue); HET = heterozygous cells (yellow); MUT = *JAK2^V617F^* mutant cells (red); NA = not assigned (grey). **f**, Relative fold change of mutant fraction in MkP and EP clusters compared to HSPCs across all patient samples. Samples for which more than one technical replicate was performed are shown separately. Statistical significance was determined by Fisher test followed by Bonferroni multiple hypothesis correction. Family-wise error rate (FWER) indicated as *P_adj_;* n.s. = not significant (*P_adj_* > 0.05). **g**, Relative fold change of mutant fraction in MkP and EP clusters compared to HSPC cluster for the fedratinib treated sample Pt-07. Statistical significance was determined by Fisher test followed by Bonferroni multiple hypothesis correction. Family-wise error rate (FWER) indicated as *P_adj_;* n.s. = not significant (*P_adj_* > 0.05). **h**, UMAP illustrating semi-supervised pseudotime estimation for hematopoietic lineages: megakaryocytic (left panel), and monocyte (right panel). **i**, Fraction of mutant cells along megakaryocyte or monocyte pseudotime for untreated or ruxolitinib-treated samples. Pseudotime was divided in 8 quantiles; each point represents the mean fraction of mutant cells, error bars indicate standard error, lines indicate the fit and shadowed areas represent the 95% confidence interval of the generalized additive model. The fraction of cells belonging to the cluster specified by color (HSPC = hematopoietic stem and progenitor cell; MPP = multipotent progenitor; MEP = megakaryocyte erythrocyte progenitor; MkP = megakaryocyte progenitor; LMPP = lymphoid-myeloid pluripotent progenitor; GMP = granulocyte monocyte progenitor; CD14 Mono = CD14^+^ monocyte progenitors; CD16 Mono = CD16^+^ monocytes) within the indicated pseudotime quantile is shown (bottom panel).

**Extended Data Fig. 5.**
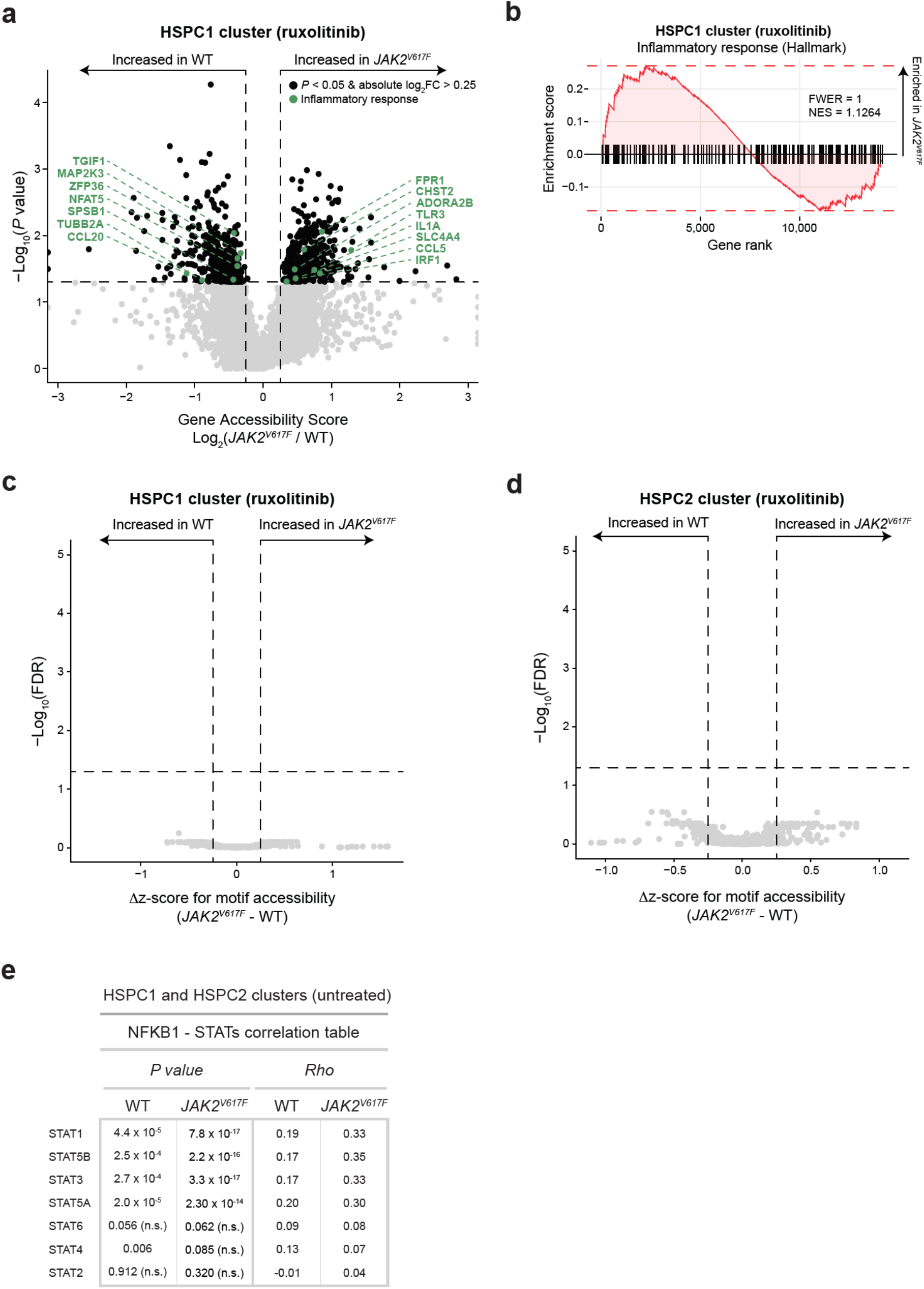
Differential gene and transcription factor motif accessibility in ruxolitinib-treated HSPCs. **a**, Volcano plot illustrating differential gene accessibility scores between wildtype (n = 55 cells) and mutant (n = 87 cells) cells within the HSPC1 cluster of ruxolitinib-treated patients (n = 3). Horizontal dotted line represents *P* = 0.05; vertical dotted lines represent absolute log_2_FC > 0.25. Genes involved in the inflammatory response pathway (Hallmark M5932) are highlighted in green. Linear mixture model (LMM) followed by likelihood ratio test. **b**, Pre-ranked gene set enrichment of genes within the Inflammatory response pathway for wildtype vs *JAK2^V617F^* HSPC1 gene accessibility scores in ruxolitinib-treated samples (Bonferroni multiple hypothesis correction; FWER = family wise error rate; NES = normalized enrichment score; Hallmark pathway M5932). **c**, Volcano plot of differential motif accessibility scores between wildtype and mutant cells within the HSPC1 cluster from patients treated with ruxolitinib JAK inhibitor therapy. Horizontal dotted lines indicate FDR = 0.25, vertical dotted lines indicate absolute Δz-score > 0.25. Linear mixture model followed by likelihood ratio test and Benjamini-Hochberg correction. **d**, Volcano plot of differential motif accessibility scores between wildtype and mutant cells within the HSPC2 cluster from patients treated with ruxolitinib JAK inhibitor therapy. Horizontal dotted lines indicate FDR = 0.25, vertical dotted lines indicate absolute Δz-score > 0.25. Linear mixture model followed by likelihood ratio test and Benjamini-Hochberg correction. **e**, Table showing the correlation values and Rho goodness of fit values for HSPCs (HSPC1 and HSPC2 clusters combined) in untreated MF samples; Spearman correlation test.

**Extended Data Fig. 6.**
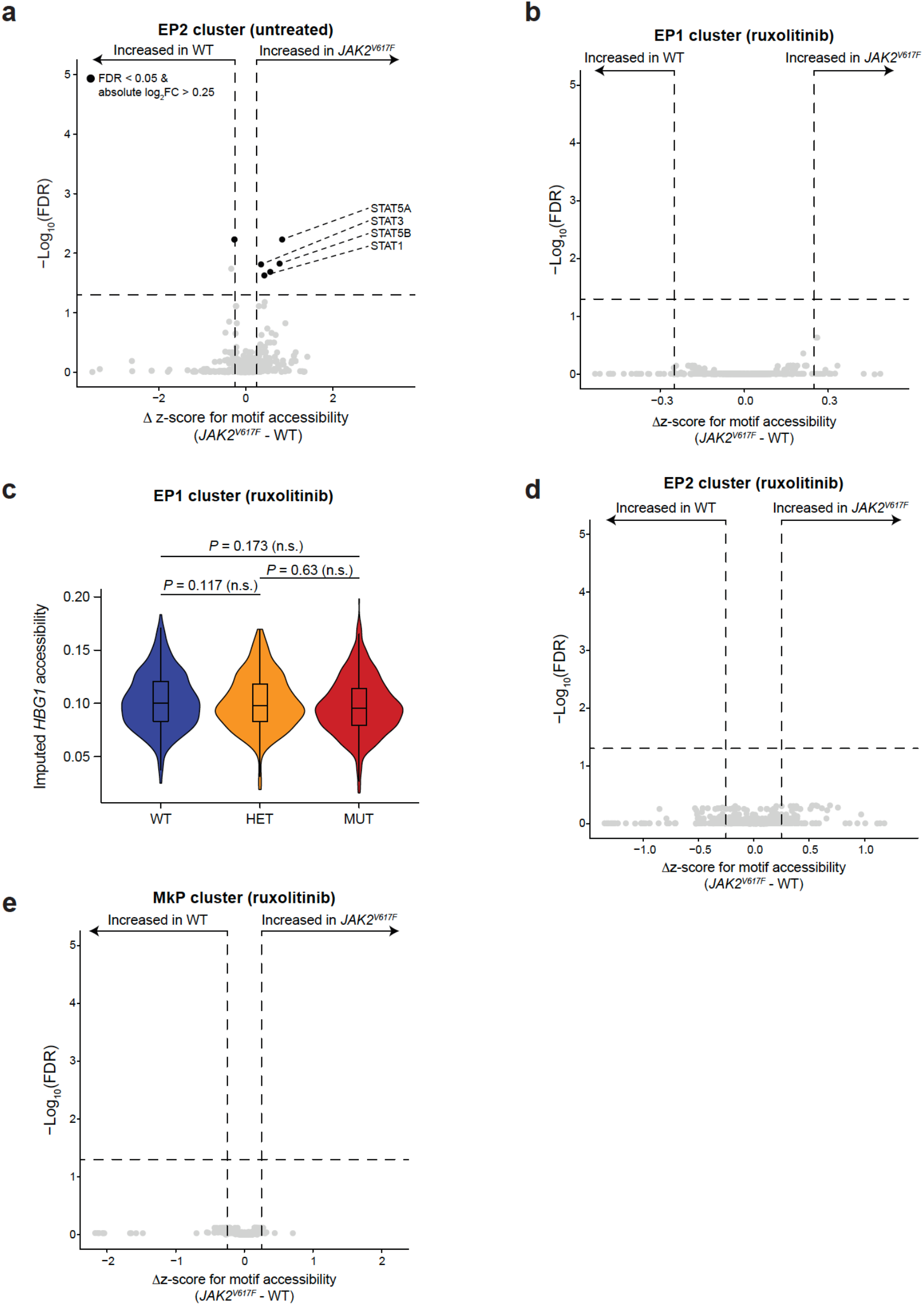
Differential gene and transcription factor motif accessibility in EPs and MkPs with and without ruxolitinib treatment. **a**, Differentially accessible transcription factor motifs between wildtype and mutant cells within the EP2 cluster of untreated MF patients (n = 4). For all panels unless stated otherwise, horizontal dotted line represents FDR = 0.05; vertical dotted lines represent absolute Δz-score> 0.25. Linear mixture model (LMM) followed by Benjamini-Hochberg correction. **b**, Differentially accessible transcription factor motifs between wildtype and mutant cells within the EP1 cluster of ruxolitinib-treated MF patients (n = 3). **c**, Quantification of the imputed gene accessibility value for either homozygous wildtype, heterozygous, or homozygous mutant cells in ruxolitinib-treated MF patients (n = 3) from the EP1 (n = 2,065 cells) cluster (LMM followed by likelihood ratio test). **d**, Differentially accessible transcription factor motifs between wildtype and mutant cells within the EP2 cluster of ruxolitinib-treated MF patients (n = 3). **e**, Differentially accessible transcription factor motifs between wildtype and mutant cells within the MkP cluster (n = 1,437 cells) of ruxolitinib-treated MF patients (n = 3).

**Extended Data Fig. 7.**
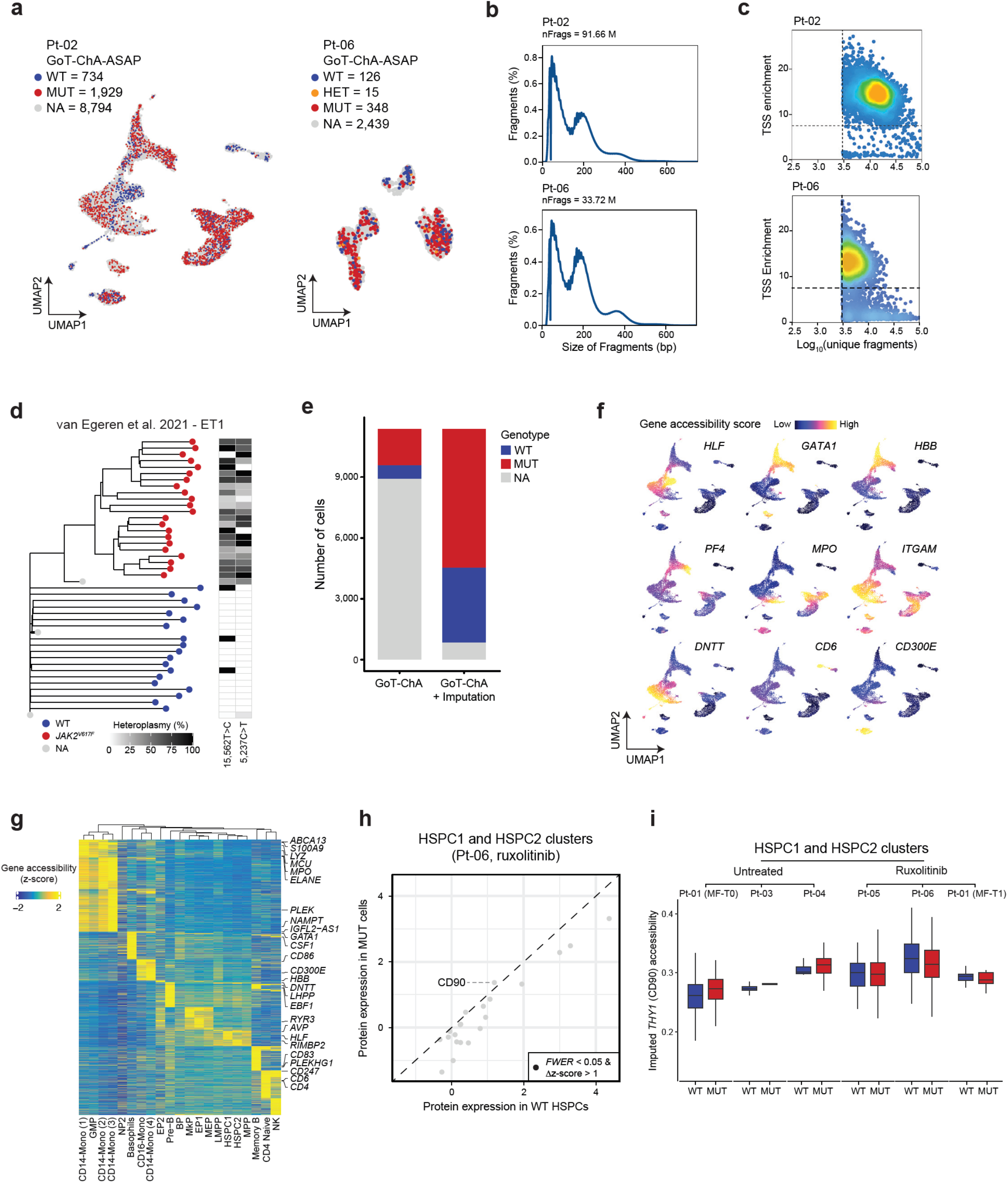
Quality control and clustering of GoT-ChA-ASAP samples. **a**, Chromatin-based uniform manifold approximation and projection (UMAPs) colored by GoT-ChA-assigned *JAK2^V617^* genotypes for Pt-02 and Pt-06 as homozygous wildtype (WT; blue), homozygous mutant (MUT; red), heterozygous (HET; yellow), and not assignable (NA; grey) prior to mitochondrial imputation. Cell numbers for each genotype are indicated per patient sample. **b**, Library fragment size distribution for samples Pt-02 and Pt-06, illustrating characteristic periodicity due to nucleosome positioning. **c**, Scatter plot showing the number of unique nuclear fragments versus TSS enrichment for samples Pt-02 and Pt-06, with dotted lines indicating the thresholds used for identification of high-quality cells (TSS enrichment score > 8 and logio [unique fragments] > 3.5). **d**, Lineage tree of HSPCs from a patient with essential thrombocythemia (ET)^31^. The phylogeny was built from 21,430 clonal SNVs detected within the single-cell expanded clones across the whole genome using CellPhy^169^. Terminal nodes are colored based on the mutation status of the *JAK2* gene. Cell heteroplasmies for two mitochondrial mutations are shown in the heatmap at the right. **e**, Total number of cells pre and post mitochondrial imputation of genotyping assignments for sample Pt-02. **f**, UMAPs showing the gene accessibility scores of canonical marker genes used to identify progenitor cell type for each cluster for sample Pt-02. **g**, Heat map of differential gene accessibility scores (FDR < 0.05 and log_2_FC > 1) across all progenitor subclusters for Pt-02. **h**, Differential protein expression between wildtype and mutant cells within HSPC cluster of ruxolitinib-treated sample Pt-06. Black dots represent differentially expressed (family wise error rate [FWER] < 0.05 and absolute Δz-score > 1; Wilcoxon rank sum test followed by Bonferroni correction) cell surface proteins. **i**, CD90 (*THY1* gene) chromatin accessibility scores of HSPCs (HSPC1 and HSPC2 clusters) per patient for either untreated or ruxolitinib-treated patients included in Fig. 5k.

## REFERENCES

1. Corces, M. R. et al. Lineage-specific and single-cell chromatin accessibility charts human hematopoiesis and leukemia evolution. Nat Genet 48, 1193–1203 (2016).

2. Buenrostro, J. D. et al. Single-cell chromatin accessibility reveals principles of regulatory variation. Nature 523, 486–490 (2015).

3. Ma, S. et al. Chromatin Potential Identified by Shared Single-Cell Profiling of RNA and Chromatin. Cell 183, 1103–1116.e20 (2020).

4. Rodrigues, C. P., Shvedunova, M. & Akhtar, A. Epigenetic Regulators as the Gatekeepers of Hematopoiesis. Trends in Genetics 37, 125–142 (2021).

5. Nam, A. S. et al. Single-cell multi-omics of human clonal hematopoiesis reveals that DNMT3A R882 mutations perturb early progenitor states through selective hypomethylation. bioRxiv 2022.01.14.476225 (2022) doi:10.1101/2022.01.14.476225.

6. Izzo, F. et al. DNA methylation disruption reshapes the hematopoietic differentiation landscape. Nature Genetics 52, 378–387 (2020).

7. Mullally, A. et al. Physiological Jak2V617F Expression Causes a Lethal Myeloproliferative Neoplasm with Differential Effects on Hematopoietic Stem and Progenitor Cells. Cancer Cell 17, 584–596 (2010).

8. Dupont, S. et al. The JAK2 617V>F mutation triggers erythropoietin hypersensitivity and terminal erythroid amplification in primary cells from patients with polycythemia vera. Blood 110, 1013–1021 (2007).

9. Gerritsen, M. et al. RUNX1 mutations enhance self-renewal and block granulocytic differentiation in human in vitro models and primary AMLs. Blood Advances 3, 320–332 (2019).

10. Renneville, A. et al. Cooperating gene mutations in acute myeloid leukemia: a review of the literature. Leukemia 22, 915–931 (2008).

11. Jaiswal, S. et al. Age-Related Clonal Hematopoiesis Associated with Adverse Outcomes. New England Journal of Medicine 371, 2488–2498 (2014).

12. Abelson, S. et al. Prediction of acute myeloid leukaemia risk in healthy individuals. Nature 559, 400–404 (2018).

13. Desai, P. et al. Somatic mutations precede acute myeloid leukemia years before diagnosis. Nature Medicine 24, 1015–1023 (2018).

14. Bick, A. G. et al. Inherited causes of clonal haematopoiesis in 97,691 whole genomes. Nature 1–7 (2020) doi:10.1038/s41586-020-2819-2.

15. Baxter, E. J. et al. Acquired mutation of the tyrosine kinase JAK2 in human myeloproliferative disorders. The Lancet 365, 1054–1061 (2005).

16. Levine, R. L. et al. Activating mutation in the tyrosine kinase JAK2 in polycythemia vera, essential thrombocythemia, and myeloid metaplasia with myelofibrosis. Cancer Cell 7, 387–397 (2005).

17. James, C. et al. A unique clonal JAK2 mutation leading to constitutive signalling causes polycythaemia vera. Nature 434, 1144–1148 (2005).

18. Kralovics, R. et al. A Gain-of-Function Mutation of JAK2 in Myeloproliferative Disorders. New England Journal of Medicine 352, 1779–1790 (2005).

19. Koschmieder, S. et al. Myeloproliferative neoplasms and inflammation: whether to target the malignant clone or the inflammatory process or both. Leukemia 30, 1018–1024 (2016).

20. Mondet, J., Hussein, K. & Mossuz, P. Circulating Cytokine Levels as Markers of Inflammation in Philadelphia Negative Myeloproliferative Neoplasms: Diagnostic and Prognostic Interest. Mediators of Inflammation 2015, e670580 (2015).

21. Tefferi, A. et al. Circulating Interleukin (IL)-8, IL-2R, IL-12, and IL-15 Levels Are Independently Prognostic in Primary Myelofibrosis: A Comprehensive Cytokine Profiling Study. JCO 29, 1356–1363 (2011).

22. Marty, C. et al. Myeloproliferative neoplasm induced by constitutive expression of JAK2V617F in knock-in mice. Blood 116, 783–787 (2010).

23. Jamieson, C. H. M. et al. The JAK2 V617F mutation occurs in hematopoietic stem cells in polycythemia vera and predisposes toward erythroid differentiation. PNAS 103, 6224–6229 (2006).

24. Giustacchini, A. et al. Single-cell transcriptomics uncovers distinct molecular signatures of stem cells in chronic myeloid leukemia. Nature Medicine 23, 692–702 (2017).

25. Rodriguez-Meira, A. et al. Unravelling Intratumoral Heterogeneity through High-Sensitivity Single-Cell Mutational Analysis and Parallel RNA Sequencing. Molecular Cell 73, 1292–1305.e8 (2019).

26. Kleppe, M. et al. JAK-STAT Pathway Activation in Malignant and Nonmalignant Cells Contributes to MPN Pathogenesis and Therapeutic Response. Cancer Discovery 5, 316–331 (2015).

27. Rodriguez-Meira, A., O’Sullivan, J., Rahman, H. & Mead, A. J. TARGET-Seq: A Protocol for High-Sensitivity Single-Cell Mutational Analysis and Parallel RNA Sequencing. STAR Protocols 1, 100125 (2020).

28. van Galen, P. et al. Single-Cell RNA-Seq Reveals AML Hierarchies Relevant to Disease Progression and Immunity. Cell 176, 1265–1281.e24 (2019).

29. Nam, A. S. et al. Somatic mutations and cell identity linked by Genotyping of Transcriptomes. Nature 571, 355–360 (2019).

30. Petti, A. A. et al. A general approach for detecting expressed mutations in AML cells using single cell RNA-sequencing. Nature Communications 10, 1–16 (2019).

31. Van Egeren, D. et al. Reconstructing the Lineage Histories and Differentiation Trajectories of Individual Cancer Cells in Myeloproliferative Neoplasms. Cell Stem Cell 28, 514–523.e9 (2021).

32. Van Egeren, D. et al. Transcriptional differences between JAK2-V617F and wild-type bone marrow cells in patients with myeloproliferative neoplasms. Experimental Hematology (2021) doi:10.1016/j.exphem.2021.12.364.

33. Rodriguez-Meira, A. et al. Deciphering TP53 mutant Cancer Evolution with Single-Cell Multi-Omics. bioRxiv 2022.03.28.485984 (2022) doi:10.1101/2022.03.28.485984.

34. Morita, K. et al. Clonal evolution of acute myeloid leukemia revealed by high-throughput single-cell genomics. Nat Commun 11, 5327 (2020).

35. Miles, L. A. et al. Single-cell mutation analysis of clonal evolution in myeloid malignancies. Nature 587, 477–482 (2020).

36. Satpathy, A. T. et al. Massively parallel single-cell chromatin landscapes of human immune cell development and intratumoral T cell exhaustion. Nature Biotechnology 37, 925–936 (2019).

37. Lareau, C. A. et al. Massively parallel single-cell mitochondrial DNA genotyping and chromatin profiling. Nature Biotechnology 1–11 (2020) doi:10.1038/s41587-020-0645-6.

38. Mimitou, E. P. et al. Scalable, multimodal profiling of chromatin accessibility, gene expression and protein levels in single cells. Nat Biotechnol 1–13 (2021) doi:10.1038/s41587-021-00927-2.

39. Verstovsek, S. et al. A Double-Blind, Placebo-Controlled Trial of Ruxolitinib for Myelofibrosis. New England Journal of Medicine 366, 799–807 (2012).

40. Schieber, M., Crispino, J. D. & Stein, B. Myelofibrosis in 2019: moving beyond JAK2 inhibition. Blood Cancer J. 9, 1–11 (2019).

41. Pardanani, A. & Tefferi, A. Definition and management of ruxolitinib treatment failure in myelofibrosis. Blood Cancer Journal 4, e268–e268 (2014).

42. Cervantes, F. et al. Three-year efficacy, safety, and survival findings from COMFORT-II, a phase 3 study comparing ruxolitinib with best available therapy for myelofibrosis. Blood 122, 4047–4053 (2013).

43. Wernig, G. et al. Unifying mechanism for different fibrotic diseases. Proceedings of the National Academy of Sciences 114, 4757–4762 (2017).

44. Ishikawa, G. et al. Shared and Tissue-Specific Expression Signatures between Bone Marrow from Primary Myelofibrosis and Essential Thrombocythemia. Experimental Hematology 79, 16–25.e3 (2019).

45. Cui, L. et al. Activation of JUN in fibroblasts promotes pro-fibrotic programme and modulates protective immunity. Nat Commun 11, 2795 (2020).

46. Mulè, M. P., Martins, A. J. & Tsang, J. S. Normalizing and denoising protein expression data from droplet-based single cell profiling. Nat Commun 13, 2099 (2022).

47. Quentmeier, H., MacLeod, R. a. F., Zaborski, M. & Drexler, H. G. JAK2 V617F tyrosine kinase mutation in cell lines derived from myeloproliferative disorders. Leukemia 20, 471–476 (2006).

48. Dawson, M. A. et al. JAK2 phosphorylates histone H3Y41 and excludes HP1α from chromatin. Nature 461, 819–822 (2009).

49. MacKinnon, R. N. et al. Genome organization and the role of centromeres in evolution of the orythroleukaemia cell line HEL. Evolution, Medicine, and Public Health 2013, 225–240 (2013).

50. Ghandi, M. et al. Next-generation characterization of the Cancer Cell Line Encyclopedia. Nature 569, 503–508 (2019).

51. DepMap 22Q1 Public. (2022) doi:10.6084/m9.figshare.19139906.v1.

52. Stuart, T., Srivastava, A., Madad, S., Lareau, C. A. & Satija, R. Singlecell chromatin state analysis with Signac. Nat Methods 18, 1333–1341 (2021).

53. Ong, S.-M. et al. A Novel, Five-Marker Alternative to CD16-CD14 Gating to Identify the Three Human Monocyte Subsets. Frontiers in Immunology 10, (2019).

54. Hao, Y. et al. Dictionary learning for integrative, multimodal, and scalable single-cell analysis. bioRxiv 2022.02.24.481684 (2022) doi:10.1101/2022.02.24.481684.

55. Mustjoki, S. et al. JAK2V617F mutation and spontaneous megakaryocytic or erythroid colony formation in patients with essential thrombocythaemia (ET) or polycythaemia vera (PV). Leukemia Research 33, 54–59 (2009).

56. Verstovsek, S. et al. Safety and Efficacy of INCB018424, a JAK1 and JAK2 Inhibitor, in Myelofibrosis. New England Journal of Medicine 363, 1117–1127 (2010).

57. Fisher, D. A. C., Fowles, J. S., Zhou, A. & Oh, S. T. Inflammatory Pathophysiology as a Contributor to Myeloproliferative Neoplasms. Frontiers in Immunology 12, (2021).

58. Vukotić, M. et al. Inhibition of proinflammatory signaling impairs fibrosis of bone marrow mesenchymal stromal cells in myeloproliferative neoplasms. Exp Mol Med 54, 273–284 (2022).

59. Dunbar, A. et al. CXCL8/CXCR2 signaling mediates bone marrow fibrosis and represents a therapeutic target in myelofibrosis. bioRxiv 2021.12.08.471791 (2021) doi:10.1101/2021.12.08.471791.

60. Cai, Z. et al. Inhibition of Inflammatory Signaling in Tet2 Mutant Preleukemic Cells Mitigates Stress-Induced Abnormalities and Clonal Hematopoiesis. Cell Stem Cell 23, 833–849.e5 (2018).

61. Hormaechea-Agulla, D. et al. Chronic infection drives Dnmt3a-loss-of-function clonal hematopoiesis via IFNγ signaling. Cell Stem Cell 28, 1428–1442.e6 (2021).

62. Avagyan, S. et al. Resistance to inflammation underlies enhanced fitness in clonal hematopoiesis. Science 374, 768–772 (2021).

63. Caiado, F., Pietras, E. M. & Manz, M. G. Inflammation as a regulator of hematopoietic stem cell function in disease, aging, and clonal selection. Journal of Experimental Medicine 218, e20201541 (2021).

64. Hayden, M. S. & Ghosh, S. Regulation of NF-κB by TNF family cytokines. Semin Immunol 26, 253–266 (2014).

65. Mossadegh-Keller, N. et al. M-CSF instructs myeloid lineage fate in single haematopoietic stem cells. Nature 497, 239–243 (2013).

66. Yao, H. et al. Corepressor Rcor1 is essential for murine erythropoiesis. Blood 123, 3175–3184 (2014).

67. Li, Y. S. et al. The expression of monocyte chemotactic protein (MCP-1) in human vascular endothelium in vitro and in vivo. Mol Cell Biochem 126, 61–68 (1993).

68. Cominal, J. G. et al. Bone Marrow Soluble Mediator Signatures of Patients With Philadelphia Chromosome-Negative Myeloproliferative Neoplasms. Front Oncol 11, 665037 (2021).

69. Meijer, B., Gearry, R. B. & Day, A. S. The role of S100A12 as a systemic marker of inflammation. Int J Inflam 2012, 907078 (2012).

70. Schep, A. N., Wu, B., Buenrostro, J. D. & Greenleaf, W. J. chromVAR: inferring transcription-factor-associated accessibility from single-cell epigenomic data. Nature Methods 14, 975–978 (2017).

71. Thurman, R. E. et al. The accessible chromatin landscape of the human genome. Nature 489, 75–82 (2012).

72. Kleppe, M. et al. Dual Targeting of Oncogenic Activation and Inflammatory Signaling Increases Therapeutic Efficacy in Myeloproliferative Neoplasms. Cancer Cell 33, 29–43.e7 (2018).

73. Fujioka, S. et al. NF-κB and AP-1 Connection: Mechanism of NF-κB-Dependent Regulation of AP-1 Activity. Molecular and Cellular Biology 24, 7806–7819 (2004).

74. Ji, Z., He, L., Regev, A. & Struhl, K. Inflammatory regulatory network mediated by the joint action of NF-kB, STAT3, and AP-1 factors is involved in many human cancers. Proceedings of the National Academy of Sciences 116, 9453–9462 (2019).

75. Grivennikov, S. I. & Karin, M. Dangerous liaisons: STAT3 and NF-κB collaboration and crosstalk in cancer. Cytokine & Growth Factor Reviews 21, 11–19 (2010).

76. Chen, E. et al. Distinct Clinical Phenotypes Associated with JAK2V617F Reflect Differential STAT1 Signaling. Cancer Cell 18, 524–535 (2010).

77. Chiu, S. K. et al. A novel role for Lyl1 in primitive erythropoiesis. Development 145, dev162990 (2018).

78. Souroullas, G. P., Salmon, J. M., Sablitzky, F., Curtis, D. J. & Goodell, M. A. Adult hematopoietic stem and progenitor cells require either Lyl1 or Scl for survival. Cell Stem Cell 4, 180–186 (2009).

79. Carmichael, C. L. et al. The EMT modulator SNAI1 contributes to AML pathogenesis via its interaction with LSD1. Blood 136, 957–973 (2020).

80. in ‘t Hout, F. E. M. et al. TCF4 promotes erythroid development. Experimental Hematology 69, 17–21.e1 (2019).

81. Chan, S. S.-K. et al. Mesp1 Patterns Mesoderm into Cardiac, Hematopoietic, or Skeletal Myogenic Progenitors in a ContextDependent Manner. Cell Stem Cell 12, 587–601 (2013).

82. Bungartz, G., Land, H., Scadden, D. T. & Emerson, S. G. NF-Y is necessary for hematopoietic stem cell proliferation and survival. Blood 119, 1380–1389 (2012).

83. Zhu, J., Zhang, Y., Joe, G. J., Pompetti, R. & Emerson, S. G. NF-Ya activates multiple hematopoietic stem cell (HSC) regulatory genes and promotes HSC self-renewal. Proceedings of the National Academy of Sciences 102, 11728–11733 (2005).

84. Lu, Z. et al. Polycomb Group Protein YY1 Is an Essential Regulator of Hematopoietic Stem Cell Quiescence. Cell Reports 22, 1545–1559 (2018).

85. Völkel, S. et al. Zinc Finger Independent Genome-Wide Binding of Sp2 Potentiates Recruitment of Histone-Fold Protein Nf-y Distinguishing It from Sp1 and Sp3. PLOS Genetics 11, e1005102 (2015).

86. Völkel, S. et al. Transcription factor Sp2 potentiates binding of the TALE homeoproteins Pbx1:Prep1 and the histone-fold domain protein Nf-y to composite genomic sites. Journal of Biological Chemistry 293, 19250–19262 (2018).

87. Zhao, Q. et al. Combined Id1 and Id3 Deletion Leads to Severe Erythropoietic Disturbances. PLOS ONE 11, e0154480 (2016).

88. Hansson, A., Zetterblad, J., van Duren, C., Axelson, H. & Jönsson, J.-I. The Lim-only protein LMO2 acts as a positive regulator of erythroid differentiation. Biochemical and Biophysical Research Communications 364, 675–681 (2007).

89. Wadman, I. A. et al. The LIM-only protein Lmo2 is a bridging molecule assembling an erythroid, DNA-binding complex which includes the TAL1, E47, GATA-1 and Ldb1/NLI proteins. The EMBO Journal 16, 3145–3157 (1997).

90. Stanulović, V. S., Cauchy, P., Assi, S. A. & Hoogenkamp, M. LMO2 is required for TAL1 DNA binding activity and initiation of definitive haematopoiesis at the haemangioblast stage. Nucleic Acids Research 45, 9874–9888 (2017).

91. Bianchi, E. et al. c-myb supports erythropoiesis through the transactivation of KLF1 and LMO2 expression. Blood 116, e99–e110 (2010).

92. Wang, J. et al. Interplay between the EMT transcription factors ZEB1 and ZEB2 regulates hematopoietic stem and progenitor cell differentiation and hematopoietic lineage fidelity. PLOS Biology 19, e3001394 (2021).

93. Robb, L. et al. The scl gene product is required for the generation of all hematopoietic lineages in the adult mouse. EMBOJ 15, 4123–4129 (1996).

94. Yokomizo, T. et al. Runx1 is involved in primitive erythropoiesis in the mouse. Blood 111, 4075–4080 (2008).

95. Willcockson, M. A. et al. Runx1 promotes murine erythroid progenitor proliferation and inhibits differentiation by preventing Pu.1 downregulation. Proc Natl Acad Sci U S A 116, 17841–17847 (2019).

96. Kim, M. Y. et al. Mbd2-CP2c loop drives adult-type globin gene expression and definitive erythropoiesis. Nucleic Acids Res 46, 4933–4949 (2018).

97. Starnes, L. M. et al. NFI-A directs the fate of hematopoietic progenitors to the erythroid or granulocytic lineage and controls β-globin and G-CSF receptor expression. Blood 114, 1753–1763 (2009).

98. Chen, L. et al. Transcriptional diversity during lineage commitment of human blood progenitors. Science 345, 1251033 (2014).

99. Wang, C. Q. et al. Cbfb deficiency results in differentiation blocks and stem/progenitor cell expansion in hematopoiesis. Leukemia 29, 753–757 (2015).

100. de Bruijn, M. & Dzierzak, E. Runx transcription factors in the development and function of the definitive hematopoietic system. Blood 129, 2061–2069 (2017).

101. Tahmasebi, S. et al. Control of embryonic stem cell self-renewal and differentiation via coordinated alternative splicing and translation of YY2. Proceedings of the National Academy of Sciences 113, 12360–12367 (2016).

102. Gao, J., Chen, Y.-H. & Peterson, L. C. GATA family transcriptional factors: emerging suspects in hematologic disorders. Experimental Hematology & Oncology 4, 28 (2015).

103. Crispino, J. D. & Horwitz, M. S. GATA factor mutations in hematologic disease. Blood 129, 2103–2110 (2017).

104. Grass, J. A. et al. GATA-1-dependent transcriptional repression of GATA-2 via disruption of positive autoregulation and domain-wide chromatin remodeling. Proceedings of the National Academy of Sciences 100, 8811–8816 (2003).

105. Komorowska, K. et al. Hepatic Leukemia Factor Maintains Quiescence of Hematopoietic Stem Cells and Protects the Stem Cell Pool during Regeneration. Cell Reports 21, 3514–3523 (2017).

106. Budi, E. H., Schaub, J. R., Decaris, M., Turner, S. & Derynck, R. TGF-β as a driver of fibrosis: physiological roles and therapeutic opportunities. The Journal of Pathology 254, 358–373 (2021).

107. Bock, O. et al. Bone Morphogenetic Proteins Are Overexpressed in the Bone Marrow of Primary Myelofibrosis and Are Apparently Induced by Fibrogenic Cytokines. The American Journal of Pathology 172, 951–960 (2008).

108. Ciaffoni, F. et al. Activation of non-canonical TGF-β1 signaling indicates an autoimmune mechanism for bone marrow fibrosis in primary myelofibrosis. Blood Cells Mol Dis 54, 234–241 (2015).

109. Martyré, M. C. et al. Transforming growth factor-β and megakaryocytes in the pathogenesis of idiopathic myelofibrosis. British Journal of Haematology 88, 9–16 (1994).

110. Dunbar, A. et al. Jak2V617F Reversible Activation Shows an Essential Requirement for Jak2V617F in Myeloproliferative Neoplasms. bioRxiv (2022).

111. Grebien, F. et al. Stat5 activation enables erythropoiesis in the absence of EpoR and Jak2. Blood 111, 4511–4522 (2008).

112. Damen, J. E. et al. Tyrosine 343 in the erythropoietin receptor positively regulates erythropoietin-induced cell proliferation and Stat5 activation. EMBO J 14, 5557–5568 (1995).

113. Wang, V. E. H., Schmidt, T., Chen, J., Sharp, P. A. & Tantin, D. Embryonic Lethality, Decreased Erythropoiesis, and Defective Octamer-Dependent Promoter Activation in Oct-1-Deficient Mice. Molecular and Cellular Biology 24, 1022–1032 (2004).

114. Gao, X. et al. Thyroid hormone receptor beta and NCOA4 regulate terminal erythrocyte differentiation. Proceedings of the National Academy of Sciences 114, 10107–10112 (2017).

115. Kendrick, T. S. et al. Erythroid defects in TRα-/-mice. Blood 111, 3245–3248 (2008).

116. Pina, C. et al. Single-Cell Network Analysis Identifies DDIT3 as a Nodal Lineage Regulator in Hematopoiesis. Cell Reports 11, 1503–1510 (2015).

117. Burda, P., Laslo, P. & Stopka, T. The role of PU.1 and GATA-1 transcription factors during normal and leukemogenic hematopoiesis. Leukemia 24, 1249–1257 (2010).

118. Will, B. et al. Minimal PU.1 reduction induces a preleukemic state and promotes development of acute myeloid leukemia. Nat Med 21, 1172–1181 (2015).

119. Blaybel, R., Théoleyre, O., Douablin, A. & Baklouti, F. Downregulation of the Spi-1/PU.1 oncogene induces the expression of TRIM10/HERF1, a key factor required for terminal erythroid cell differentiation and survival. Cell Res 18, 834–845 (2008).

120. Basak, A. & Sankaran, V. G. Regulation of the fetal hemoglobin silencing factor BCL11A. Annals of the New York Academy of Sciences 1368, 25–30 (2016).

121. Lamsfus-Calle, A. et al. Comparative targeting analysis of KLF1, BCL11A, and HBG1/2 in CD34+ HSPCs by CRISPR/Cas9 for the induction of fetal hemoglobin. Sci Rep 10, 10133 (2020).

122. Bauer, D. E. et al. An Erythroid Enhancer of BCL11A Subject to Genetic Variation Determines Fetal Hemoglobin Level. Science 342, 253–257 (2013).

123. Pliner, H. A. et al. Cicero Predicts cis-Regulatory DNA Interactions from Single-Cell Chromatin Accessibility Data. Molecular Cell 71, 858–871.e8 (2018).

124. Jackson, J. D., Petrykowska, H., Philipsen, S., Miller, W. & Hardison, R. Role of DNA sequences outside the cores of DNase hypersensitive sites (HSs) in functions of the beta-globin locus control region. Domain opening and synergism between HS2 and HS3. J Biol Chem 271, 11871–11878 (1996).

125. Mendek-Czajkowska, E. et al. Hemoglobin F in primary myelofibrosis and in myelodysplasia. Clinical & Laboratory Haematology 25, 289–292 (2003).

126. Rinaldi, C. R., Rinaldi, P., Pane, F., Camera, A. & Rinaldi, C. Acquired Hb H disease associated with elevated Hb F level in patient affected by primary myelofibrosis. Ann Hematol 89, 827–828 (2010).

127. Hoffman, R. et al. Fetal hemoglobin in polycythemia vera: cellular distribution in 50 unselected patients. Blood 53, 1148–1155 (1979).

128. Ciurea, S. O. et al. Pivotal contributions of megakaryocytes to the biology of idiopathic myelofibrosis. Blood 110, 986–993 (2007).

129. Jeremy Wen, Q. et al. Targeting megakaryocytic-induced fibrosis in myeloproliferative neoplasms by AURKA inhibition. Nat Med 21, 1473–1480 (2015).

130. Castro-Malaspina, H., Rabellino, E. M., Yen, A., Nachman, R. L. & Moore, M. A. S. Human Megakaryocyte Stimulation of Proliferation of Bone Marrow Fibroblasts. Blood 57, 781–787 (1981).

131. Baum, C. M., Weissman, I. L., Tsukamoto, A. S., Buckle, A. M. & Peault, B. Isolation of a candidate human hematopoietic stem-cell population. Proc Natl Acad Sci U S A 89, 2804–2808 (1992).

132. Tong, J. et al. Hematopoietic Stem Cell Heterogeneity Is Linked to the Initiation and Therapeutic Response of Myeloproliferative Neoplasms. Cell Stem Cell 28, 502–513.e6 (2021).

133. McGranahan, N. & Swanton, C. Clonal Heterogeneity and Tumor Evolution: Past, Present, and the Future. Cell 168, 613–628 (2017).

134. Mustjoki, S. & Young, N. S. Somatic Mutations in “Benign” Disease. New England Journal of Medicine 384, 2039–2052 (2021).

135. Martincorena, I. et al. Somatic mutant clones colonize the human esophagus with age. Science 362, 911–917 (2018).

136. Martincorena, I. et al. High burden and pervasive positive selection of somatic mutations in normal human skin. Science 348, 880–886 (2015).

137. Yizhak, K. et al. RNA sequence analysis reveals macroscopic somatic clonal expansion across normal tissues. Science 364, eaaw0726 (2019).

138. Yokoyama, A. et al. Age-related remodelling of oesophageal epithelia by mutated cancer drivers. Nature 565, 312–317 (2019).

139. Yoshida, K. et al. Tobacco smoking and somatic mutations in human bronchial epithelium. Nature 578, 266–272 (2020).

140. Kakiuchi, N. & Ogawa, S. Clonal expansion in non-cancer tissues. Nature Reviews Cancer 1–18 (2021) doi:10.1038/s41568-021-00335-3.

141. Kakiuchi, N. et al. Frequent mutations that converge on the NFKBIZ pathway in ulcerative colitis. Nature 577, 260–265 (2020).

142. Lee-Six, H. et al. The landscape of somatic mutation in normal colorectal epithelial cells. Nature 574, 532–537 (2019).

143. Harrison, C. et al. JAK Inhibition with Ruxolitinib versus Best Available Therapy for Myelofibrosis. New England Journal of Medicine 366, 787–798 (2012).

144. Patel, K. P. et al. Correlation of mutation profile and response in patients with myelofibrosis treated with ruxolitinib. Blood 126, 790–797 (2015).

145. Pacilli, A. et al. Mutation landscape in patients with myelofibrosis receiving ruxolitinib or hydroxyurea. Blood Cancer Journal 8, 1–10 (2018).

146. Newberry, K. J. et al. Clonal evolution and outcomes in myelofibrosis after ruxolitinib discontinuation. Blood 130, 1125–1131 (2017).

147. Jayavelu, A. K. et al. Splicing factor YBX1 mediates persistence of JAK2-mutated neoplasms. Nature 1–7 (2020) doi:10.1038/s41586-020-2968-3.

148. Tefferi, A. et al. The clinical phenotype of wild-type, heterozygous, and homozygous JAK2V617F in polycythemia vera. Cancer 106, 631–635 (2006).

149. Passamonti, F. et al. A prospective study of 338 patients with polycythemia vera: the impact of JAK2 (V617F) allele burden and leukocytosis on fibrotic or leukemic disease transformation and vascular complications. Leukemia 24, 1574–1579 (2010).

150. Vannucchi, A. M. et al. Clinical profile of homozygous JAK2 617V>F mutation in patients with polycythemia vera or essential thrombocythemia. Blood 110, 840–846 (2007).

151. Vannucchi, A. M., Pieri, L. & Guglielmelli, P. JAK2 allele burden in the myeloproliferative neoplasms: effects on phenotype, prognosis and change with treatment. Therapeutic Advances in Hematology 2, 21–32 (2011).

152. Campbell, P. J. et al. Mutation of JAK2 in the myeloproliferative disorders: timing, clonality studies, cytogenetic associations, and role in leukemic transformation. Blood 108, 3548–3555 (2006).

153. Akada, H. et al. Conditional expression of heterozygous or homozygous Jak2V617F from its endogenous promoter induces a polycythemia vera-like disease. Blood 115, 3589–3597 (2010).

154. Xu, W. et al. ISSAAC-seq enables sensitive and flexible multimodal profiling of chromatin accessibility and gene expression in single cells. bioRxiv 2022.01.16.476488 (2022) doi:10.1101/2022.01.16.476488.

155. Stuart, T. et al. Nanobody-tethered transposition allows for multifactorial chromatin profiling at single-cell resolution. bioRxiv 2022.03.08.483436 (2022) doi:10.1101/2022.03.08.483436.

156. Bartosovic, M., Kabbe, M. & Castelo-Branco, G. Single-cell CUT&Tag profiles histone modifications and transcription factors in complex tissues. Nat Biotechnol 39, 825–835 (2021).

157. Granja, J. M. et al. ArchR is a scalable software package for integrative single-cell chromatin accessibility analysis. Nat Genet 53, 403–411 (2021).

158. Mulè, M. P., Martins, A. J. & Tsang, J. S. Normalizing and denoising protein expression data from droplet-based single cell profiling. Nat Commun 13, 2099 (2022).

159. Stoeckius, M. et al. Cell Hashing with barcoded antibodies enables multiplexing and doublet detection for single cell genomics. Genome Biol 19, 224 (2018).

160. Hao, Y. et al. Integrated analysis of multimodal single-cell data. Cell 184, 3573–3587.e29 (2021).

161. Stuart, T. et al. Comprehensive Integration of Single-Cell Data. Cell 177, 1888–1902.e21 (2019).

162. Bray, N. L., Pimentel, H., Melsted, P. & Pachter, L. Near-optimal probabilistic RNA-seq quantification. Nat Biotechnol 34, 525–527 (2016).

163. Melsted, P. et al. Modular, efficient and constant-memory singlecell RNA-seq preprocessing. Nat Biotechnol 39, 813–818 (2021).

164. Gehring, J., Hwee Park, J., Chen, S., Thomson, M. & Pachter, L. Highly multiplexed single-cell RNA-seq by DNA oligonucleotide tagging of cellular proteins. Nat Biotechnol 38, 35–38 (2020).

165. Ludwig, L. S. et al. Lineage Tracing in Humans Enabled by Mitochondrial Mutations and Single-Cell Genomics. Cell 176, 1325–1339.e22 (2019).

166. Svetnik, V. et al. Random Forest: A Classification and Regression Tool for Compound Classification and QSAR Modeling. J. Chem. Inf. Comput. Sci. 43, 1947–1958 (2003).

167. Plummer, N. W. et al. Expanding the power of recombinase-based labeling to uncover cellular diversity. Development 142, 4385–4393 (2015).

168. Ruzankina, Y. et al. Deletion of the Developmentally Essential Gene ATR in Adult Mice Leads to Age-Related Phenotypes and Stem Cell Loss. Cell Stem Cell 1, 113–126 (2007).

169. Kozlov, A., Alves, J. M., Stamatakis, A. & Posada, D. CellPhy: accurate and fast probabilistic inference of single-cell phylogenies from scDNA-seq data. Genome Biology 23, 37 (2022).

